# Extreme sex chromosome differentiation, likely driven by inversion, contrasts with mitochondrial paraphyly between species of crowned sparrows

**DOI:** 10.1101/2022.08.19.504329

**Authors:** Quinn McCallum, Kenneth Askelson, Finola Fogarty, Libby Natola, Ellen Nikelski, Andrew Huang, Darren Irwin

## Abstract

Sympatric species pairs provide researchers with the opportunity to study patterns of genomic differentiation during the late stages of speciation and to identify the genomic regions underlying reproductive isolation. The Golden-crowned Sparrow (*Zonotrichia atricapilla*) and the White-crowned Sparrow (*Zonotrichia leucophrys*) are broadly sympatric songbirds found in western North America. These sister species are phenotypically differentiated and largely reproductively isolated despite possessing similar mitochondrial genomes, likely due to recent mitochondrial introgression. We used a genotyping-by-sequencing (GBS) approach to determine the structure of nuclear genomic differentiation between these species and also between two hybridizing subspecies of *Z. leucophrys*, across more than 45,000 single nucleotide polymorphisms (SNPs). The two *Z. leucophrys* subspecies showed moderate levels of relative differentiation, as well as patterns consistent with a history of recurrent selection in both ancestral and daughter populations. *Z. leucophrys* and *Z. atricapilla* show high levels of relative differentiation and strong heterogeneity in the level of differentiation among different chromosomal regions, with a large portion of the Z chromosome showing highly elevated differentiation. Patterns of relative and absolute differentiation and linkage disequilibrium suggest a large inversion on the Z chromosome, with inversion haplotypes that segregate between *Z. atricapilla* and *Z. leucophrys*. While mitochondrial DNA differentiation is often emphasized in studies of speciation, differentiation between these *Zonotrichia* sparrows appears to have occurred first in the Z chromosome and secondarily in autosomes, followed by mitochondrial introgression. This putative inversion has implications for reproductive isolation between these species and adds to a growing body of evidence for the importance of inversions and the Z chromosome in speciation.

## Introduction

Understanding how the genome evolves during speciation is of great interest to evolutionary biologists. Often, the nuclear genomes of diverging populations show highly heterogeneous patterns of differentiation (e.g., Irwin et al., 2018; Toews, Taylor, et al., 2016). Specific regions of the genome that show elevated differentiation can reveal genetic or chromosomal drivers of speciation. These differentiation peaks, also referred to as “islands of differentiation”, are typically the result of divergent selection and genetic hitchhiking (Cruickshank & Hahn, 2014; Irwin et al., 2016) or the presence of a locus that resists introgression (Cruickshank & Hahn, 2014). Thus, by comparing relative differentiation (*F*_ST_), absolute differentiation (*π*_Between_, also known as *D*_XY_), and nucleotide diversity (*π*_Within_), we can identify historical selection and introgression in particular genomic regions. At the same time, genome-wide comparisons of these statistics allow inference of the evolutionary processes that occurred during speciation.

Models typically invoked to explain peaks in differentiation include divergence with gene flow, divergence in allopatry, recurrent selection in ancestral and daughter populations, or introgression and selective sweeps followed by differentiation (Cruickshank & Hahn, 2014; Irwin et al., 2016, 2018). Empirical studies often find support for multiple patterns in different areas of the genome, and all these processes can act together to shape patterns of genomic differentiation. Wild birds have served as important natural systems to study genomic differentiation, and have helped shape our understanding of speciation (reviewed in Toews, Campagna, et al., 2016). For example, many North American Passerine superspecies share a common biogeographic history of repeated isolation and secondary contact due to glacial cycles throughout the Pleistocene (Weir & Schluter, 2004). Some of these superspecies show a negative correlation between *F*_ST_ and *π*_Between_, suggesting a history of repeated bouts of differentiation followed by hybridization and introgression, during periods of isolation and secondary contact caused by glacial cycles over the course of speciation (Irwin et al., 2018).

In birds, sex chromosomes (Z and W) often play an outsized role in speciation, as numerous studies report increased *F*_ST_ and reduced *π*_Within_ across the Z chromosome relative to autosomes (reviewed by Irwin, 2018; Mank et al., 2009; Presgraves, 2018). The effective population size of the avian Z chromosome is three quarters that of the autosomes, thus reduced nucleotide diversity across the Z chromosome is expected. However, many observed values of *π*_Z_/*π*_A_ fall far below neutral expectations, implicating selection (Irwin, 2018). Differentiation can also be promoted by structural variants in the genome that reduce recombination, such as chromosomal inversions (Ortiz-Barrientos et al., 2016). Crossover events between inversion haplotypes cause large deletions and duplications, often resulting in inviable gametes by heterozygotes. This effectively prevents recombination between inversion haplotypes, preventing gene flow in these regions and allowing for the accumulation of differentiation (Noor et al., 2001; Ortiz-Barrientos et al., 2016; Rieseberg, 2001). Chromosomal inversions are thus often implicated in local adaptation (e.g., Lowry & Willis, 2010; Todesco et al., 2020a) and divergence between species (e.g. Hooper et al., 2019; Noor et al., 2001). Finally, narrow peaks of increased *F*_ST_ can be used to identify specific genes that have diverged between populations, providing valuable insights into genes and phenotypes that underlie reproductive isolation (reviewed in Toews, Campagna, et al., 2016).

In contrast to the nuclear genome, differentiation across the length of the mitochondrial genome is usually less variable. The mitochondrial genome lacks recombination and is thus inherited as a single locus (Ballard & Whitlock, 2004). Historically, the mitochondrial genome was considered a neutral marker and was used extensively in early molecular phylogenetics (e.g. Zink et al., 1991). However, recent scholarship suggests that the mitochondrial genome and its products are subject to selection (e.g., Hill 2019). Furthermore, hybridization can result in introgression of the mitochondrial genome across species boundaries (Wang et al. 2021) and an introgressed mitochondrial haplotype may then be subject to a selective sweep (e.g., Irwin et al. 2009; Taylor et al., 2021). Such a phenomenon results in discordance between the phylogenetic reconstructions based on mitochondrial and nuclear loci—a pattern observed in numerous avian clades (reviewed in Toews & Brelsford 2012). Here, we investigated the genomic landscape of differentiation between one such pair of birds which show deep divergence in the nuclear genome, but paraphyly of the mitochondrial genome (Taylor et al. 2021).

The Golden-crowned Sparrows (*Zonotrichia atricapilla*) and White-crowned Sparrows (*Zonotrichia leucophrys*) are an excellent model for studying genomic differentiation at the end stages of speciation, despite suggestions of recent gene flow between groups. These sister species show marked phenotypic differences (Zink 1982) and are easily distinguished by the color and pattern of their crowns (Fig. 1A). Interestingly, *Z. leucophrys* shows geographic variation between five recognized subspecies which differ slightly in bill colour, plumage, and migratory behaviour (Chilton et al., 2020). Previous phylogenetic work has found that the widely distributed *Z. l. leucophrys, Z. l. orianthra*, and *Z. l. gambelii* subspecies form a separate clade from *Z. l. pugetensis* and *Z. l. nuttalli* of the Pacific Coast (Taylor et al., 2021). In contrast, *Z. atricapilla* is restricted to the Pacific Coast of North America by the Rocky Mountains and shows no apparent geographic variation in phenotype (Norment et al., 2020). Notably, *Z. atricapilla* is sympatric over much of its breeding range with the *gambelii* subspecies of *Z. leucophrys* (Weckstein et al. 2001; Fig. 1A). Despite their extensive sympatry and similar ecologies, very few putative hybrids have been reported between these species, suggesting that reproductive isolation is nearly complete (Miller, 1940; Morton & Mewaldt, 1960).

**Figure 1.**
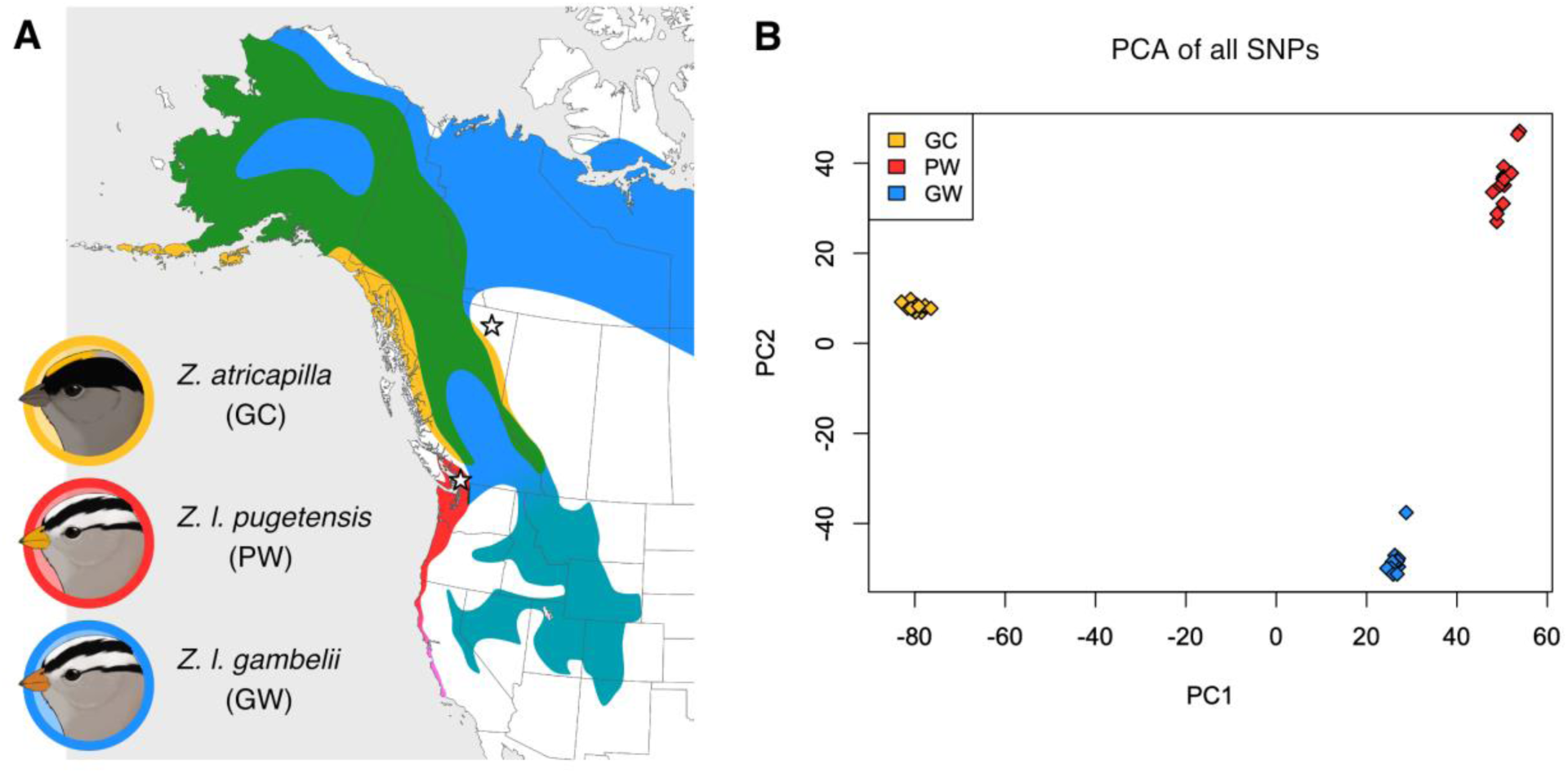
(A) Map of the breeding ranges of the Golden-crowned Sparrow (*Zonotrichia atricapilla* in yellow), the Pacific White-crowned Sparrows (*Zonotrichia leucophrys pugetensis* in red; *Zonotrichia leucophrys nuttallii* in magenta), and the Eastern White-crowned Sparrow (*Zonotrichia leucophrys gambelii* in blue; *Zonotrichia leucophrys oriantha* in teal). Regions of sympatry between *Z. atricapilla* and *Z. l. gambelii* are shown in green. Species ranges based on Chilton et al. (2020) and Norment et al. (2020). Stars represent the locations at which samples were collected. (B) Principal component analysis of 45,986 SNPs across the genome. Yellow represents *Z. atricapilla* (GC) individuals, blue represents *Z. l. gambelii* (GW), and red represents *Z. l. pugetensis* (PW). PC1 explains 24.4% of the variation in the dataset. PC2 explains 9.28% of the variation in the dataset.

While the rate of hybridization between *Z. atricapilla* and *Z. leucophrys* is apparently low, there is strong evidence for extensive mitochondrial introgression between the two taxa (Taylor et al., 2021; Weckstein et al., 2001). Early molecular phylogenies of *Zonotrichia* found remarkably little sequence differentiation between the mitochondrial genomes (mtDNA) of these species (Zink et al., 1991; Zink & Blackwell, 1996) but marked differences in allozyme sequences and morphology (Zink & Blackwell, 1996). This low mtDNA differentiation was further investigated by Weckstein et al. (2001) and later by Taylor et al. (2021), who found that the two species shared mitochondrial haplotypes and were not reciprocally monophyletic based on mtDNA. However, the species showed deep divergence in a phylogeny built from nuclear loci (Taylor et al., 2021). This discordance between the nuclear and mitochondrial phylogenies is strong evidence for mitochondrial introgression between the species, possibly followed by a selective sweep for the introgressed *Z. leucophrys* mitochondrial genome in *Z. atricapilla* (Taylor et al., 2021). However, previous work on this system has not explored whether the high degree of nuclear differentiation between these species is genome-wide or driven by particular regions. Taken together, the low rate of hybridization across a region of broad sympatry suggests that reproductive isolation is nearly complete between these species, but recent mitochondrial capture suggests introgression may still occur. By contrast, the coastal and interior lineages of *Z. leucophrys* are parapatric, and secondary contact between *Z. l. pugetensis* and *Z. l. gambelii* is apparently very recent (Hunn & Beaudette, 2014). However, previous studies have detected gene flow between these groups, suggesting they are not reproductively isolated (Taylor et al., 2021). Comparing between *Z. atricapilla* and *Z. leucophrya* and between the two lineages of *Z. leucophrys* allows for the examination of patterns of genomic differentiation between pairs of taxa in the same clade that show different levels of reproductive isolation.

In 2018, two putative *Z. atricapilla* x *leucophrys* hybrids were captured and banded at the Iona Island Observatory (IIBO) in Richmond, British Columbia, during spring migration monitoring efforts (Fig. S1). The following spring we took genetic samples from all *Z. atricapilla* and *Z. leucophrys* captured at IIBO, and used Genotyping by Sequencing (GBS) to survey the landscape of genomic differentiation between the two species, as well as between two distinct lineages of *Z. leucophrys.* The goals of this study were to (1) identify regions of the nuclear genome driving the overall pattern of high nuclear differentiation observed by Taylor et al. (2021) and (2) compare the patterns of genomic differentiation observed between pairs of taxa that show differing levels of divergence. Given the phylogenetic relationship of these taxa, we predicted to find higher overall differentiation and broader genomic islands of differentiation between *Z. atricapilla* and *Z. leucophrys* than between the two *Z. leucophrys* lineages. Additionally, we present evidence for elevated differentiation between *Z. atricapilla* and *Z. leucophrys* over a large region of the Z chromosome, consistent with an inversion that may have played an outsized role during speciation in these taxa. This research provides further evidence for the importance of the avian Z chromosome and inversions in speciation and contributes to our broader understanding of how genomic differentiation progresses despite occasional hybridization between species.

## Methods and Materials

### Sample Collection

The first round of fieldwork was conducted at the Iona Island Bird Observatory (IIBO) in Richmond, British Columbia, in Spring 2019. Birds were captured passively in mist nets and banded as part of the station’s normal migration monitoring operations. We then took 10 – 40 μL of blood from the brachial vein of each bird of the target species and stored these samples in 500 μL of Queen’s Lysis Buffer (Seutin et al., 1991). We measured wing chord, tail length, tarsus length, mass, culmen length, beak depth, and beak width. A total of 19 *Z. atricapilla* and 14 *Z. leucophrys* of indeterminate subspecies were sampled during this initial period.

The second round of field sampling focused on increasing the sample size of *Z*. *leucophrys* and was conducted between June 16 and 26, 2020 at the Vancouver Campus of the University of British Columbia (UBC). Singing birds were located, and a mist net was set up nearby. Song recordings of local *Z. leucophrys* were used to attract birds to the net. Once birds were captured, they were immediately removed, banded, sampled, and measured using the methods outlined above. We sampled an additional 16 breeding *Z. leucophrys* individuals during this collection period, all of the *pugetensis* subspecies based on their bill colour and location. Additionally, we took samples of pectoral muscle tissue from seven unprepared specimens from the freezer of the Cowan Tetrapod Collection at the Beaty Biodiversity Museum of UBC that had been salvaged from throughout BC.

### GBS Library Preparation

Genomic DNA was extracted from blood and tissue samples using a standard phenol-chloroform extraction procedure. DNA sample concentrations were measured using a Qubit fluorometer (Invitrogen), diluted to between 15 and 30 ng of DNA per *μ*L, then run on 2% agarose gels to assess DNA fragmentation. Three out of our 56 samples did not produce DNA of sufficient quality for use.

The samples were genotyped using a genotyping-by-sequencing (GBS) procedure first described by Elshire et al. (2011) with modifications described in Alcaide et al. (2014) and Geraldes et al. (2019). We prepared two GBS libraries with a 400-500 bp fragment size. The library containing samples from 2019 was sequenced on one lane of an Illumina HiSeq 4000 sequencer at the Genome Quebec Innovation Centre. The library containing samples from 2019 was sequenced on one lane of an Illumina NovaSeq 6000 SP also at the Genome Quebec Innovation Centre. Both sequencing platforms generated 150 bp paired-end reads.

The first stage of sequencing produced 290,502,080 reads from 94 individual birds, 37 of which were *Zonotrichia* sparrows collected in 2019 or prior for inclusion in this study (the others were for an unrelated study). The second stage of sequencing produced 525,322,895 reads from 88 individuals, 16 of which were sparrows collected in 2020 for this study. The disparity in sequence amount between the two plates was due to the different sequencing platforms used.

### Bioinformatics Pipeline

Reads were demultiplexed using custom scripts from Baute et al. (2016), and sequences were trimmed for quality using Trimmomatic (version 0.36; Bolger et al., 2014). The reads were then aligned to the *Taeniopygia guttata* reference genome version bTaeGut2.pat.W.v2 (Warren et al., 2010; Rhie et al., 2021; GenBank accession number GCA_008822105.2) using BWA-MEM (version 0.7.15; Li & Durbin, 2009). The resulting SAM files were converted to BAM files using Picard (version 2.23.8; Broad Institute https://broadinstitute.github.io/picard/), and single-end and paired-end BAM files were combined using SAMtools (version 1.3.2; Li et al., 2009). Single Nucleotide Polymorphisms (SNPs) were identified using the HaplotypeCaller tool in GATK (version 3.8; McKenna et al., 2010), which produced a gvcf file for each individual. Individual g.vcf files were then combined into a single vcf file using the GenotypeGVCFs tool in GATK. We filtered genomic sites using VCFtools (version 1.16; Danecek et al., 2011), first by removing indels and SNPs with more than 2 alleles. Using a custom Perl script (Owens et al., 2016), we removed sites with a mapping quality lower than 20 and a heterozygosity above 0.6. We then filtered for individuals with more than 60% missing data using VCFtools, removing three *Z. atricapilla* and three *Z. leucophrys* from the dataset. Finally, we used VCFtools to filter out SNPs missing from more than 20% of individuals and SNPs with a minimum allele frequency of less than 0.05.

A comparison of mean coverage for each individual across the sex chromosomes (W and Z) and a representative autosome (chromosome 3), revealed that coverage across the W chromosome was approximately three times higher than across the autosomes (Fig. S2). This was true for known female birds, as well as known males. Since females are heterogametic for the W chromosome and males are homogametic for the Z (Irwin, 2018), we expect coverage across the W to be roughly half that of autosomal coverage in females, and zero in males. These results indicate that many of the sequences from our short-read dataset that mapped to the Zebra Finch W chromosome cannot be from the *Zonotrichia* W chromosome. These likely represent sequences found on autosomes in *Zonotrichia*, but on the W chromosome in the Zebra finch, or repetitive elements found at high frequency in *Zonotrichia*, but infrequently in the Zebra Finch. After filtering and removal of the W sequences, a total of 45,986 SNPs were retained for analysis.

### Principal Component Analysis

To visualize the population genetic clustering of individuals and identify potential backcrossed hybrids in our sample, we performed Principal Component Analysis (PCA) on the genotype data using custom *R* (version 3.6.3; R Core Team) scripts by Irwin et al. (2016), and the package pcaMethods (version 1.78.0; Stacklies et al., 2007) to impute missing data using svdImpute. Individual PCAs were also generated for each chromosome using the same method.

### Patterns of Genomic Differentiation

Our genome-wide PCA showed three distinct clusters representing *Z. atricapilla, Z. l. pugetensis*, and *Z. l. gambelii* (see Results), which we treated as separate taxa for all future analysis performed each possible pairwise comparison (i.e., *Z. atricapilla* versus *Z. l. pugetensis*, *Z. atricapilla versus Z. l. gambelii*, and *Z. l. pugetensis* versus *Z. l. gambelii*). First, *F_ST_* was calculated for each filtered variant site, followed by genome-wide weighted mean *F*_ST_ using Weir and Cockerham’s (1984) estimation with VCFtools. To further investigate patterns of genomic differentiation, we calculated mean *F*_ST_, *π*_Between_ (absolute sequence divergence between species) and *π*_Within_ (absolute nucleotide diversity within species) across sliding windows of 10,000 bases of mapped sequence, including both variant and invariant sites, using R scripts by Irwin et al. (2018). Note that these windows include unsequenced regions, and thus represent regions of the genome roughly 500kb to 1Mb in length. We randomly selected 15 individuals of each taxon to include in this analysis to prevent bias from uneven sampling. To characterize genome-wide differentiation patterns, windowed data were compared in scatterplots of *π*_Between_ versus mean *π*_Within_, and *F*_ST_ versus *π*_Between_. We performed a Spearman’s rank correlation test for each of these comparisons.

### Comparing Z Chromosome and Autosomes

To test for differences in mean *F*_ST_, *π*_Between_, and *π*_Within_ between the Z chromosome and autosomes, we treated each 10,000 bp sliding window as a data point and performed a series of Welch’s two-sample *t*-tests. We compared mean *F*_ST_ across the Z chromosome (*F*_ST, Z_) to mean autosomal *F*_ST_ (*F*_ST, A_) in all comparisons. For comparisons of *π*_Between_ and *π*_Within_, we accounted for the increased mutation rate experienced by the Z chromosome by dividing the windowed stats from the Z chromosome by 1.1 (*π*_Between, Z_/ 1.1 = *π*_Between, Z*_; π_Within, Z_ / 1.1 = *π*_Within, Z*_), based on the calculations of Irwin (2018). We then compared mean *π*_Between, Z*_ to mean *π*_Between, A_ for each comparison and mean *π*_Within, Z*_ to mean *π*_Within, A_ for each taxon. Finally, we calculated ratios of mean F_ST, Z_ / F_ST, A_ and mean *π*_Between, Z*_/ *π*_Between, A_ for each comparison, and mean *π*_Within, Z*_ / *π*_Within, A_ for each taxon. To test for selection on the Z chromosome, we also performed a Welch’s *t*-test to compare mean *π*_Within,Z*_ to the mean of *π*_Within, A_ multiplied by 0.5625 (the lowest ratio that might be explainable without selection; Charlesworth, 2001; Irwin, 2018).

### Investigating Putative Inversions

The Z chromosome and chromosome 1A contained large blocks showing patterns of *F*_ST_, *π*_Between_, and *π*_Within_ consistent with inversions. To investigate this further, we calculated pairwise linkage disequilibrium (LD) between each biallelic variant site on chromosomes 1A and Z for each group separately, and with relevant groups pooled together, using Plink (version 1.9; Chang et al., 2015) and visualized the data in R. Linkage disequilibrium is the non-random association of alleles between two loci, thus, regions that show increased *F*_ST_ between groups are expected to show concordant increases LD (*R*^2^ close to 1) when calculated between groups. If these regions show low LD when calculated within groups, we can discount the possibility that the region intrinsically experiences reduced recombination rate, such as in centromeres. We also plotted the genotypes of individuals across these regions at all SNPs, and at all SNPs with an *F*_ST_ above 0.75 (Irwin et al., 2016).

## Results

### Three Genomic Clusters

Principal Component Analysis (PCA) revealed three distinct genetic clusters (Fig. 1B). *Z. atricapilla* and *Z. leucophrys* were highly separated along PC1, which accounts for 24.4% of the variation in the dataset. PC2, which accounts for 9.3% of the variation in the dataset, revealed two distinct genetic clusters in *Z. leucophrys.* These clusters are also observed on most of the PCA’s for individual chromosomes (Fig. S3). In the lower right of Figure 1B, one cluster was composed of birds captured at IIBO in 2019 during spring migration and four individuals collected in Fort Nelson, BC, in May of 2012. Fort Nelson is within the range of *Z. leucophrys gambelii*, and all these individuals showed the orange bill characteristic of that subspecies. The birds captured at IIBO were likely migrating to more northerly breeding grounds. In the upper right, the other cluster was composed primarily of birds caught at UBC during the breeding season, and one individual captured at IIBO. These birds all showed the yellow bill characteristic of the subspecies *Z. l. pugetensis.* The distinct clustering of the two species and the two *Z. leucophrys* subspecies is consistent with Taylor et al. (2021). In further analyses, we treated these two *Z. leucophrys* groups separately.

One individual from the *Z. l. gambelii* group fell slightly outside the main *gambelii* cluster, in the direction of the *pugetensis* cluster. This individual, a female captured at IIBO in 2019 during spring migration, fell midway between the two groups on the PCAs for chromosomes 1, 6, 8, 11, 12, and 14, and clustered with the *pugetensis* group on the PCA for the Z chromosome (Fig. S3). Taken together, this suggests that this individual is the result of recent hybridization and backcrossing between these two subspecies of *Z. leucophrys*. Three other individuals were intermediate between the *Z. leucophrys* groups on the PCAs for chromosome 1A, and another was intermediate for chromosome 8 (Fig. S3). These patterns are consistent with ongoing gene flow between *Z. l. gambelii* and *Z. l. pugetensis.* The PCAs do not reveal evidence for hybridization and/or gene flow between *Z. atricapilla* and *Z. leucophrys.*

### Larger Autosomes are Associated with Increased Differentiation and Reduced Nucleotide Diversity

Across the entire genome, weighted mean *F_ST_* was 0.273 between *Z. atricapilla* and *Z. l. gambelii*, 0.303 between *Z. atricapilla* and *Z. l. pugetensis*, and 0.141 between the two *Z. leucophrys* groups (Table 1). A total of 661 fixed sites (i.e., with *F*_ST_ = 1) were found between *Z. atricapilla* and *Z. l. gambelii*, and 657 fixed sites were found between *Z. atricapilla* and *Z. l. pugetensis*. In comparison, only 16 fixed sites were found between the two *Z. leucophrys* groups (see Fig. S4.)

**Table 1.** Summary of genome-wide weighted mean *F*_ST_ (Weir & Cockerham 1984) between pairwise comparisons of taxa.

| Weighted Mean $F_{ST}$ | *Z. atricapilla* | *Z. l. pugetensis* |
| --- | --- | --- |
| *Z. l. gambelii* | 0.273 | 0.141 |
| *Z. l. pugetensis* | 0.303 | - |

Analyses of *F*_ST_, *π*_Between_, and *π*_Within_ across 10,000 bp windows were largely consistent between the two comparisons of *Z. atricapilla* (Figs. 2A and 3A) with a *Z. leucophrys* group. These comparisons showed numerous regions of elevated *F*_ST_, particularly on larger autosomes, such as 1, 1A, 2, 3, 4 and 5, and on the Z chromosome. In general, regions of elevated *F*_ST_ extended over large stretches of the chromosome, forming blocks of high differentiation rather than narrow peaks. Both comparisons showed a weak, but statistically significant, negative correlation between *F*_ST_ and *π*_Between_ (Figs 2B and 3B; *Z. atricapilla* vs. *Z. l. gambelii*: Spearman’s rank correlation *ρ* = −0.2431, p < 2.2×10^−16^; *Z. atricapilla* vs. *Z. l. pugetensis*: Spearman’s rank correlation *ρ* = −0.2803, p < 2.2×10^−16^). However, the splines fitted to the scatterplots had a concave upward shape, suggesting that this correlation was mainly driven by low *F*_ST_ windows, while high *F*_ST_ windows were generally associated with moderate values of *π*_Between_. Both comparisons showed positive correlation between *π*_Between_ and mean *π*_Within_ (Figs. 2C and 3C; *Z. atricapilla* vs. *Z. l. gambelii*: Spearman’s rank correlation *ρ* = 0.8891, p < 2.2×10^−16^; *Z. atricapilla* vs. *Z. l. pugetensis*: Spearman’s rank correlation *ρ* = 0.9009, p < 2.2×10^−16^).

**Figure 2.**
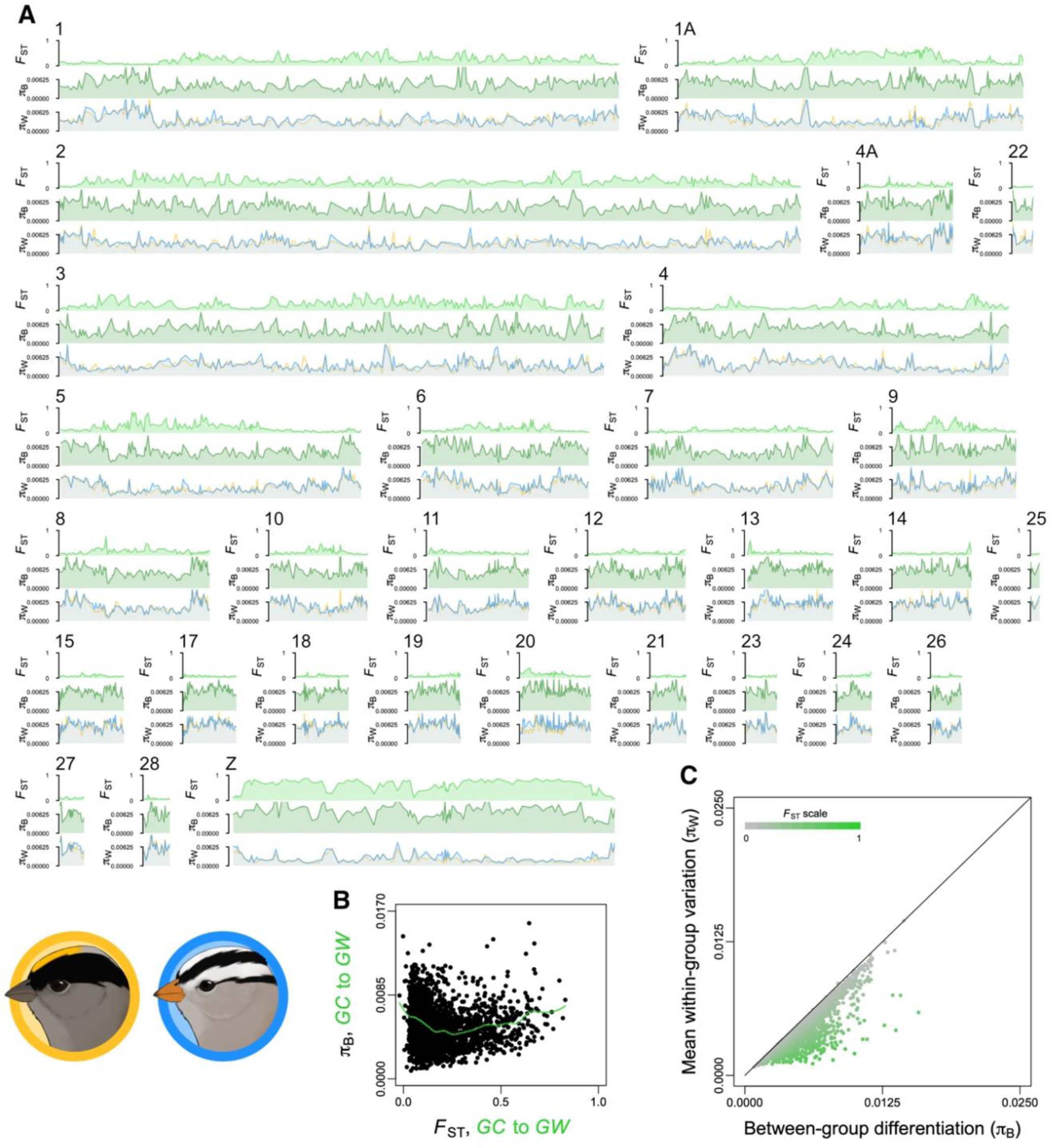
Summary of genomic differentiation and variation between *Z. atricapilla* (GC) and *Z. l. gambelii* (GW) from sliding windows containing 10,000 bp of genotyped positions. (A) Mean *F*_ST_, *π*_Between_ and *π*_Within_ for each taxon (*Z. atricapilla* in yellow and *Z. leucophrys* in blue) are shown plotted against the position for each chromosome. (B) Scatterplots in which each point represents a single 10,000 bp window depict relationships between *F*_ST_ and *π*_Between_ (Spearman’s rank correlation, *ρ* = −0.243, p < 2.2×10^−16^), and (C) between *π*_Between_ and mean *π*_Within_ (Spearman’s rank correlation *ρ* = 0.889, p < 2.2×10^−16^).

**Figure 3.**
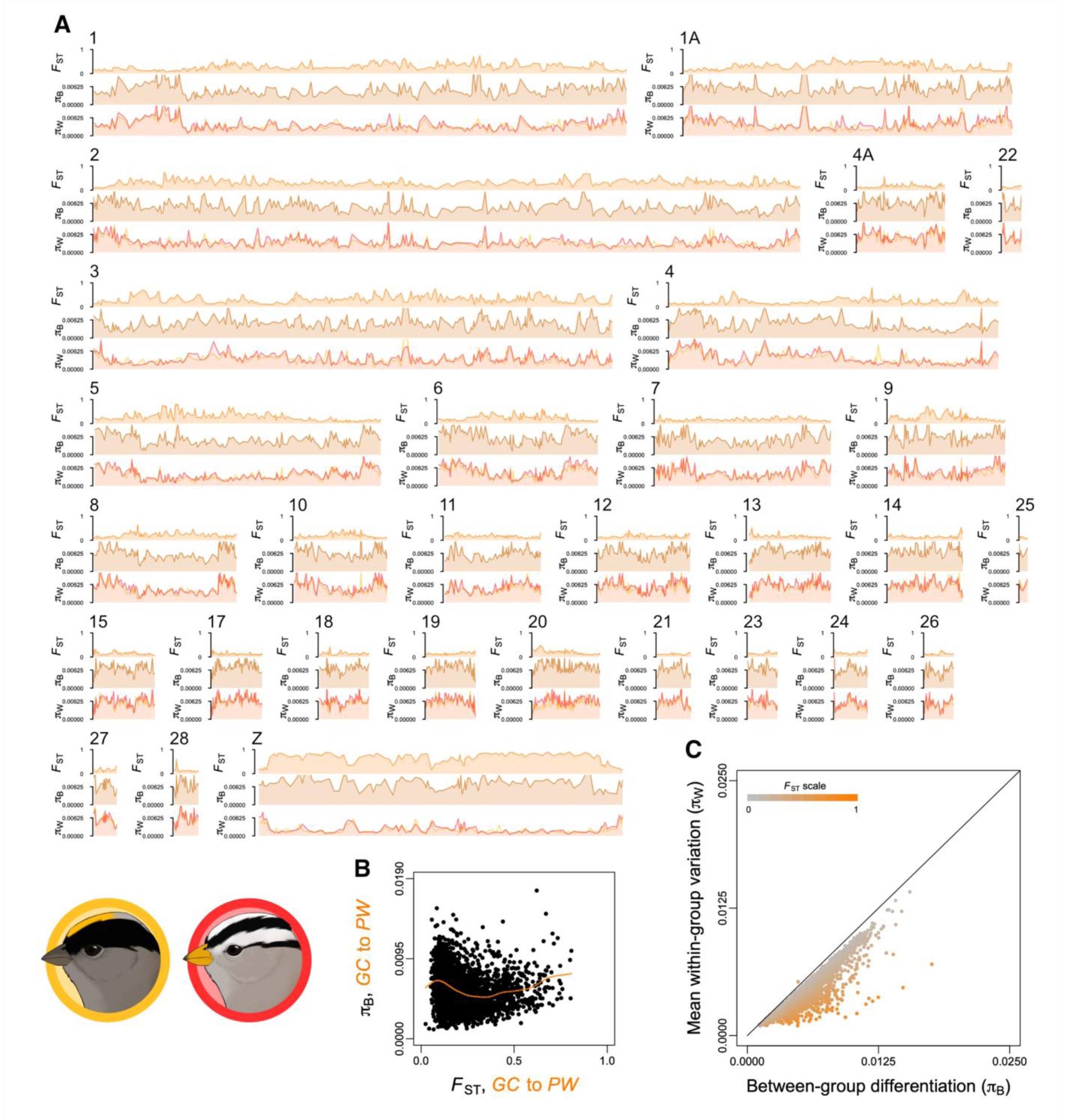
Summary of genomic differentiation and variation between *Z. atricapilla* (GC) and *Z. l. pugetensis* (PW) from sliding windows containing 10,000 bp of genotyped positions. (A) Mean *F*_ST_, *π*_Between_ and *π*_Within_ for each taxon (*Z. atricapilla* in yellow and *Z. l. pugetensis* in red) are shown plotted against the position for each chromosome. (B) Scatterplots, where each point represents a single 10,000 bp window, depict relationships between *F*_ST_ and *π*_Between_ (Spearman’s rank correlation *ρ* = −0.280, p < 2.2×10^−16^), and (C) between *π*_Between_ and mean *π*_Within_ (Spearman’s rank correlation *ρ* = 0.901, p < 2.2×10^−16^).

The windowed analysis comparing the two *Z. leucophrys* groups showed lower overall differentiation, but still with some notable regions of elevated differentiation concentrated on larger chromosomes, particularly 1A, 5 and the Z Chromosome (Fig. 4A). In this comparison, the correlation between *F*_ST_ and *π*_Between_ was also negative and stronger than between the comparisons that included *Z. atricapilla* (Fig. 4B; Spearman’s rank correlation *ρ* = −0.3183, p < 0.0001). However, once again, the highest *F*_ST_ windows showed moderate values of *π*_Between_. In this comparison, *π*_Between_ and mean *π*_Within_ showed a strong positive correlation (Fig. 4C; Spearman’s rank correlation *ρ* = 0.9752, p < 0.0001).

**Figure 4.**
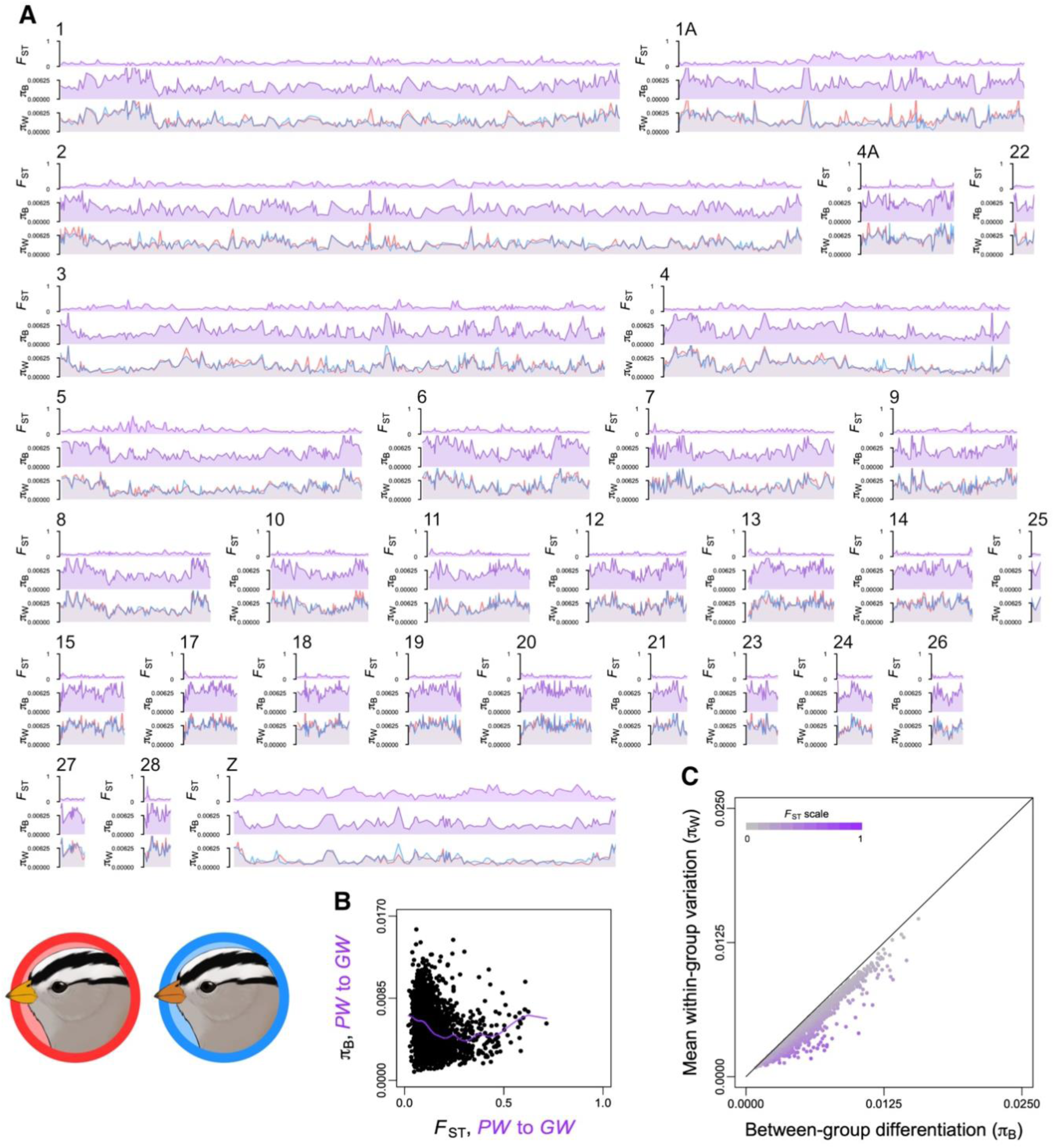
Summary of genomic differentiation and variation between *Z. l. gambelii* (GW) and *Z. l. pugetensis* (PW) from sliding windows containing 10,000 bp of genotyped positions. (A) Mean *F*_ST_, *π*_Between_ and *π*_Within_ for each taxon (*Z. l. gambelii* in blue and *Z. l. pugetensis* in red) are plotted against each chromosome’s position. (B) Scatterplots, where each point represents a single 10,000 bp window, depict relationships between *F*_ST_ and *π*_Between_ (Spearman’s rank correlation *ρ* = −0.318, p < 2.2×10^−16^), and (C) between *π*_Between_ and mean *π*_Within_ (C; Spearman’s rank correlation *ρ* = 0.975, p < 2.2×10^−16^).

In all comparisons, *F*_ST_ tended to increase with increasing chromosome length (Fig. 5). Both *π*_Between_ and *π*_Within_ decreased with increasing chromosomal size (Fig. 5). Furthermore, chromosomal PCAs for smaller chromosomes showed less distinct clustering than larger ones (Fig. S3). The PCAs for chromosomes 25 and 28 show overlap between the *Z. atricapilla* and *Z. l. gambelii* clusters, with the *Z. l. pugetensis* cluster somewhat separated from these (Fig. S3).

**Figure 5.**
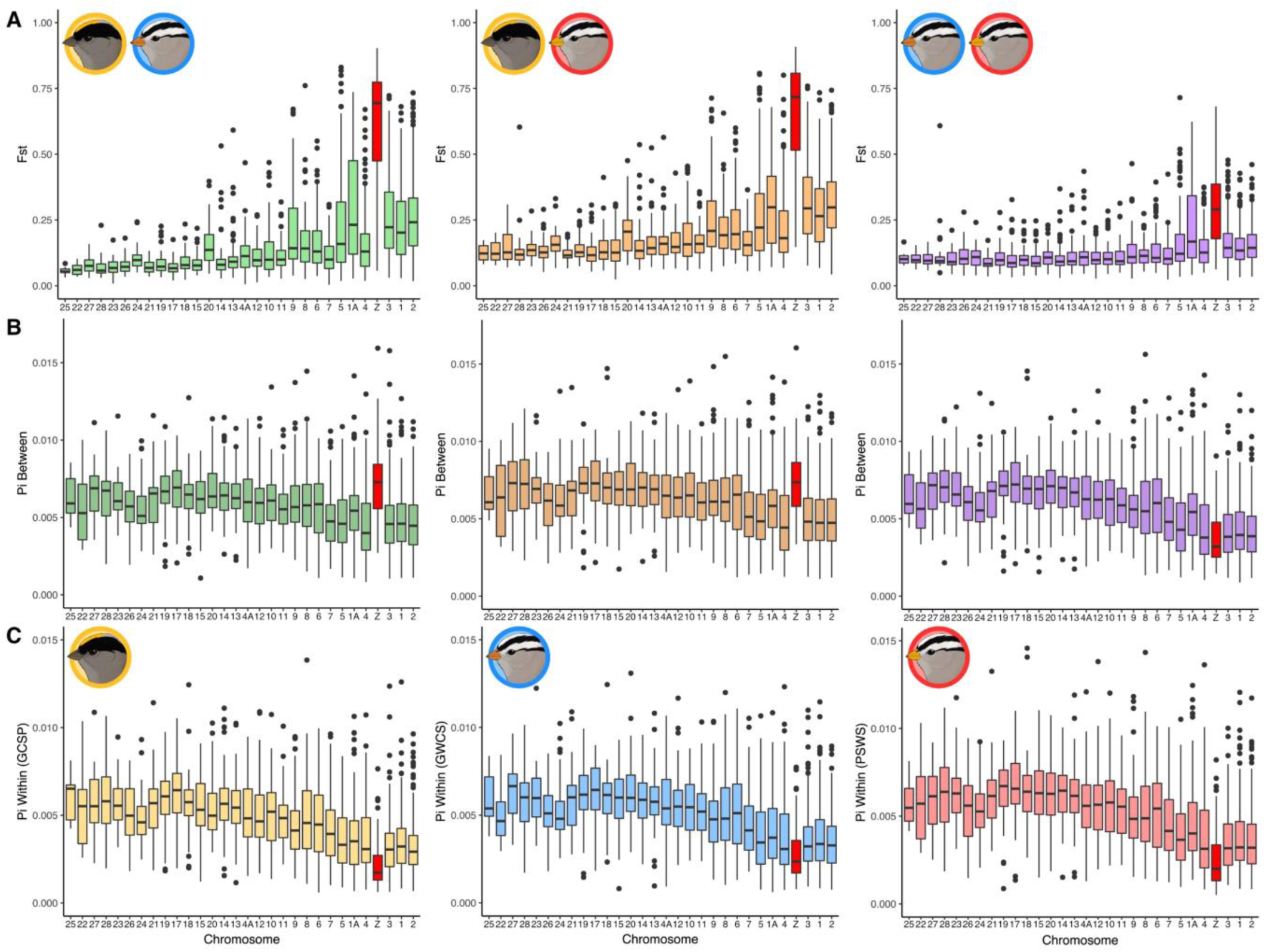
Boxplots of (A) *F*_ST_, (B) *π*_Between_ and (C) *π*_Within_ across chromosomes. Chromosomes are arranged in increasing length from left to right. In each plot, the Z chromosome is highlighted in dark red. The coloured box represents the interquartile range, and the horizontal line indicates the median. The vertical line represents values that fall within 1.5 times the interquartile range, and dots represent individual windows that fall outside this range.

### Highly Differentiated Chromosome 1A Haplotypes between *Z. leucophrys* Groups

When comparing *Z. l. gambelii* and *Z. l. pugetensis*, chromosome 1A showed high relative differentiation relative to other autosomes (Fig. 5A right panel), due to the presence of a block of high *F*_ST_ (Fig. 4A). Three *Z. l. pugetensis* individuals showed high heterozygosity across this region for *Z. l. gambelii* alleles (Fig. S4A). These individuals otherwise clustered with the remainder of the *Z. l. pugetensis* individuals on the whole genome PCA, thus these individuals do not appear to be recent hybrids. This suggests the existence of two non-recombining haplotypes in this region. Non-recombining haplotype blocks could be caused by a structural rearrangement, such as an inversion. To further investigate this possibility, we calculated and plotted LD across chromosome 1A with individuals from both subspecies, and with only individuals from each subspecies. The three highly heterozygous individuals were not included in this analysis. The LD plots showed an increase in LD concordant with the region of increased *F*_ST_ only when calculated between the two species, but not when LD is calculated within groups (Fig. S4B). This pattern is consistent with an inversion, but could also be explained by a selective sweep in a region of low recombination.

### Increased Differentiation in the Z Chromosome between *Z. atricapilla* and *Z. leucophrys*

Across all comparisons, the Z chromosome showed increased *F*_ST_ relative to the autosomes (Table 2; Welch’s two-sample *t*-tests: *Z. atricapilla* vs. *Z. l. gambelii*, *t* = 19.967, df = 123.14, p < 0.0001; *Z. atricapilla* vs. *Z. l. pugetensis*, *t* = 20.389, df = 123.03, p < 0.0001), even compared to autosomes of similar size (Fig. 5). This increase in *F*_ST,Z_ was more extreme in the comparisons involving *Z. atricapilla* than in the comparison of the *Z. leucophrys* groups. Mean *π*_Between, Z*_ was higher than mean *π*_Between, A_ in both comparisons involving *Z. atricapilla* even after adjusting for the increase in mutation rate experienced by the Z chromosome (Table 2; *Z. atricapilla* vs. *Z. l. gambelii*, *t* = 5.2402, df = 130.92, p < 0.0001; *Z. atricapilla* vs. *Z. l. pugetensis*, *t* = 4.0968, df = 134.64, p < 0.0001). Mean *π*_Between, Z*_ was reduced relative to *π*_Between, A_ between *Z. l. gambelii* and *Z. l. pugetensis* (Table 2; *t* = −14.914, df = 144.05, p < 0.0001). Finally, the Z chromosome in all three groups showed reduced *π*_Within, Z*_ compared to *π*_Within, A_, with all *π*_Within, Z*_ / *π*_Within, A_ falling below the minimum possible value expected under neutrality, 0.5625, with extreme variance in male reproductive success (Charlesworth, 2001) (Table 3; *Z. atricapilla, t* = −19.845, df = 148.9, p < 0.0001; *Z. l. gambelii, t* = −15.546, df = 141.96, p < 0.0001; *Z. l. pugetensis, t* = −17.641, df = 143.34, p < 0.0001). The PCA for the Z chromosome (Fig. S3) showed a strong separation of *Z. atricapilla* and *Z. leucophrys* along PC1, which explained 68.7% of the variation. *Z. l. gambelii* and *Z. l. pugetensis* were separated along PC2, which explained 10.6% of the variation, aside from one *Z. l. gambelii* individual which clustered with *Z. l. pugetensis.*

**Table 2.** Summary of mean *F*_ST_ across all Z chromosome windows (*F*_ST,Z_), mean F_ST_ across all autosomal windows (*F*_ST, A_), the ratio of *F*_ST,Z_ / *F*_ST, A_, mean *π*_Between_ across all Z chromosome windows (*π*_Between, Z_), mean *π*_Between, Z_ corrected for mutation rate (*π*_Between, Z*_ = *π*_Between, Z_ / 1.1), mean *π*_Between_ across all autosomal windows (*π*_Between, A_) and the ratio of *π*_Between, Z*_ / *π*_Between, A_ are given for each comparison. Results of Welch’s t-tests comparing autosomal and Z chromosome means are indicated by asterisks after each ratio. * indicates p < 0.0001.

| Comparison | $F_{ST, Z}$ | $F_{ST, A}$ | $F_{ST, Z} / F_{ST, A}$ | $\pi_{Between, Z}$ | $\pi_{Between, Z}^*$ | $\pi_{Between, A}$ | $\pi_{Between, Z}^* / \pi_{Between, A}$ |
| --- | --- | --- | --- | --- | --- | --- | --- |
| *Z. atricapilla* vs. *Z. l. gambelii* | 0.598 | 0.179 | 3.34* | 0.00707 | 0.00643 | 0.00548 | 1.17* |
| *Z. atricapilla* vs. *Z. l. pugetensis* | 0.639 | 0.230 | 2.78* | 0.00730 | 0.00663 | 0.00593 | 1.12* |
| *Z. l. pugetensis* vs. *Z. l. gambelii* | 0.298 | 0.137 | 2.17* | 0.00368 | 0.00335 | 0.00547 | 0.61* |

**Table 3.** Summary of mean *π*_Within_ across all Z chromosome windows (*π*_Within, Z_), across the Z chromosome corrected for mutation rate (*π*_Within, Z*_ = *π*_Within, Z_ / 1.1), mean *π*_Within_ across all autosomal windows (*π*_Within, A_), and the ratio of *π*_Within, Z*_ / *π*_Within, A_ for each taxon. Results of Welch’s t-tests comparing mean *π*_Within, Z*_ and *π*_Within, A_ x 0.5625 (see Irwin, 2018) are indicated by asterisks after each ratio. * indicates p < 0.01; ** indicates p < 0.001.

| Taxon | $\pi_{Within, Z}$ | $\pi_{Within, Z}^*$ | $\pi_{Within, A}$ | $\pi_{Within, Z}^* / \pi_{Within, A}$ |
| --- | --- | --- | --- | --- |
| *Z. atricapilla* | 0.00223 | 0.00202 | 0.00436 | 0.464** |
| *Z. l. gambelii* | 0.00281 | 0.00255 | 0.00467 | 0.547 |
| *Z. l. pugetensis* | 0.00244 | 0.00231 | 0.00483 | 0.477* |

Detailed plots of sliding windowed data across the Z chromosome (Fig. 6) from comparisons that include *Z. atricapilla* revealed a sudden increase in *F*_ST_ across a large block spanning the middle of the chromosome, with decreased *F*_ST_ at either end. Numerous fixed differences between the two species (SNPs with an *F*_ST_ of 1) exist within this region. In contrast, comparing the two *Z. leucophrys* groups did not show this large block of highly elevated *F*_ST_ (Fig. 6B). Furthermore, *π*_Between_ was high across the Z chromosome in the comparisons that include *Z. atricapilla* but moderate or low across this block in comparing the *Z. leucophrys* groups. In all three groups, *π*_Within_ tends to be decreased across this block relative to the ends of the chromosome.

**Figure 6.**
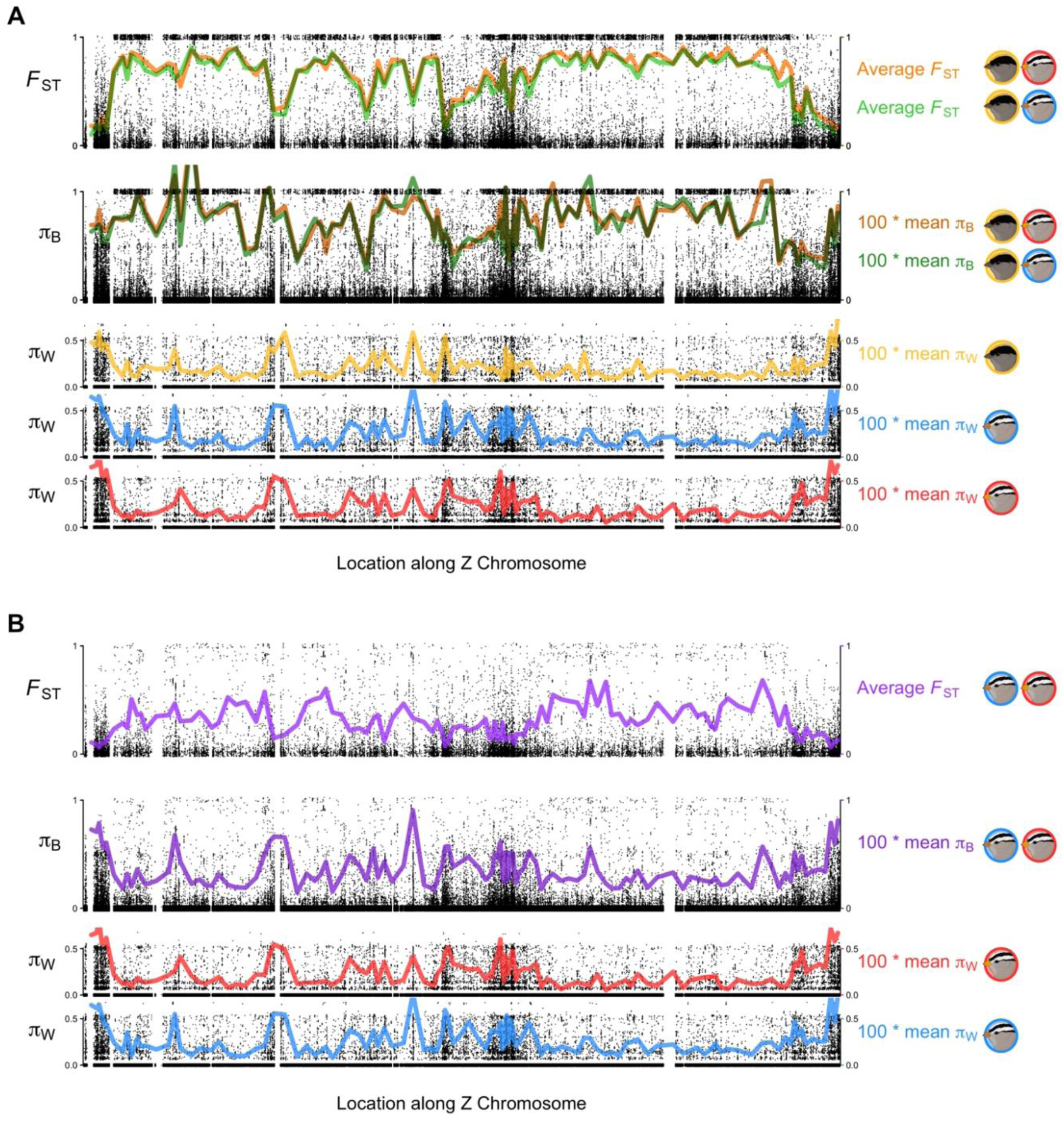
Patterns of *F*_ST_, _Between_, and _Within_ across the Z chromosome in comparisons (A) between *Z. atricapilla* and *Z. l. gambelii*, and *Z. atricapilla* and *Z. l. pugetensis* and (B) between *Z. l. gambelii* and *Z. l. pugetensis.* Black dots represent individual SNPs, while coloured lines represent average across 10,000 bp sliding windows.

The sharp increase in *F*_ST_ and fixed differences is consistent with physical linkage and reduced recombination across the Z chromosome; one possibility is that an inversion may explain this pattern. To further investigate this, we examined genotypes (Fig. 7A) and calculated LD (Fig. 7B) across the Z chromosome with individuals from both species, and with only individuals from each species. These results show a sharp increase in LD concordant with the region of increased *F*_ST_ when calculated between the two species, but not when LD is calculated within groups (Fig. 7B). Plots of the genotype of each individual at SNPs with an *F*_ST_ greater than 0.75 across the Z chromosome show two distinct haplotypes across this region of Z chromosome (Fig. 7A). One haplotype is shared by all *Z. atricapilla* individuals, while another occurs in *Z. leucophrys.*

**Figure 7.**
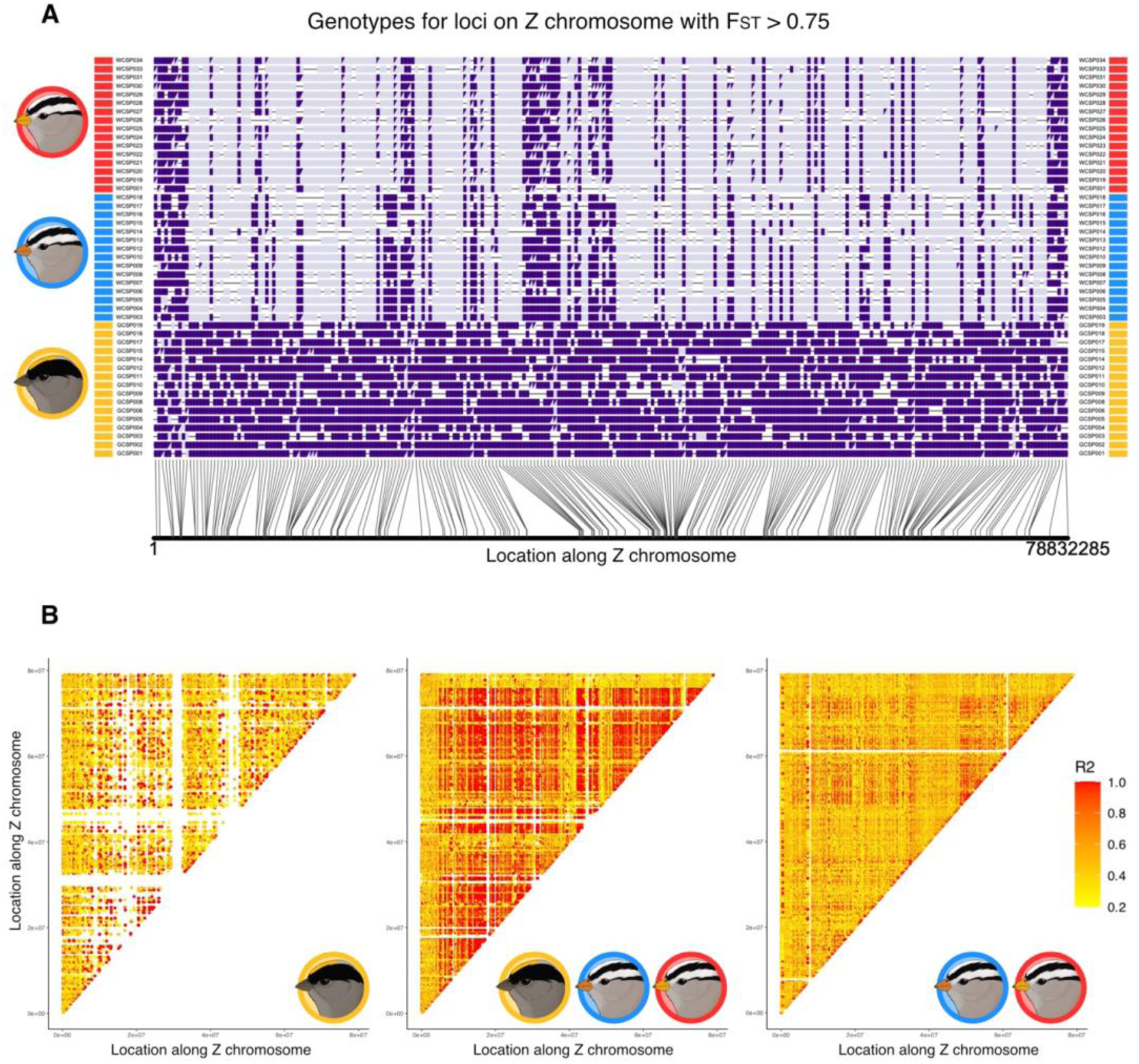
Evidence supporting a large inversion on the Z chromosome. (A) Genotypes at all biallelic SNPs that show an *F*_ST_ > 0.75 on the Z chromosome. The most common genotype in *Z. atricapilla* is plotted in dark purple, and the alternative allele is plotted in light purple. Homozygous and hemizygous loci are indicated with rectangles, and heterozygous loci are indicated with split rectangles. Each row represents an individual in the dataset, and rectangles next to rows represent the taxon of that individual, with *Z. atricapilla* in yellow, *Z. l. gambelii* in blue, and *Z. l. pugetensis* in red. Both males and females are shown. (B) Heatmaps of linkage disequilibrium (LD) across the Z chromosome when calculated between just *Z. atricapilla* (left), all three groups (middle), and just the two *Z. leucophrys* groups (right).

## Discussion

Our results show that the landscape of differentiation between the nuclear genomes of *Z. leucophrys* and *Z. atricapilla* is highly heterogeneous, with large regions of the nuclear genome showing vastly higher differentiation than that previously seen in mitochondrial DNA (Taylor et al. 2021). Differentiation peaks are concentrated on larger chromosomes, especially the Z chromosome, which appears to harbour a large inversion with haplotypes that segregate between the two species and that account for the high degree of overall nuclear differentiation observed previously (Taylor et al. 2021). This apparent Z-linked inversion may play a role in reproductive isolation between these taxa.

Genome-wide weighted mean *F*_ST_ values between *Z. atricapilla* and the *Z. leucophrys* subspecies calculated from our SNP dataset are similar to those reported by Taylor et al. (2021). These values are consistent with those typically reported in avian species pairs that show extensive reproductive isolation (e.g., Cowles & Uy, 2019; Mikkelsen & Irwin, 2021) and are much higher than other species pairs that engage in extensive hybridization (Irwin et al., 2018; Toews, Taylor, et al., 2016; Wang et al., 2020). Both comparisons show a weakly negative correlation between *F*_ST_ and *π*_Between_ for autosomal windows. However, many highly differentiated windows show low *π*_Within_, but moderate *π*_Between_ (Figs 2C and 3C). Taken together, these results suggest that autosomal divergence has primarily been driven by divergent selection, with some recurrent selection between the ancestral and contemporary species or introgression followed by differentiation, driving the slight negative correlation between *F*_ST_ and *π*_Between_ (Cruickshank & Hahn, 2014; Irwin et al., 2016, 2018). However, our sample shows no evidence of recent hybridization between these species, despite the evidence of somewhat recent mtDNA introgression. Notably, the mtDNA introgression seems to have occurred between *Z. atricapilla* and the currently allopatric *Z. l. pugetensis*, but not between *Z. atricapilla* and the currently sympatric *Z. l. gambelii* (Taylor et al., 2020; Weckstein et al., 2001). Thus, some gene flow, including mitochondrial introgression, may have occurred thousands of years ago, possibly during a recent glacial maximum when *Z. atricapilla* was sympatric with the more southerly *Z. pugetensis* (as suggested by Weckstein et al., 2001), but gene flow has since become cut off between *Z. atricapilla* and *Z. l. gambelii.*

For *Z. l. gambelii* and *Z. l. pugetensis*, patterns in *F*_ST_, *π*_Between_, and *π*_Within_ are more consistent with a history of recurrent selection (Cruickshank & Hahn, 2014), or possibly “sweep-before-differentiation” in which globally advantageous mutations spread between populations and then become differentiated (Irwin et al., 2016). This possibility is supported by our evidence of ongoing hybridization and gene flow between those taxa. Genome-wide *F*_ST_ was much lower between these taxa, which is expected given their more recent divergence. However, it is still higher than what has been reported between several boreal-breeding North American species pairs (Irwin et al., 2018). In this comparison, the correlation across autosomal windows between *F*_ST_ and *π*_Between_ is negative and stronger than in the comparisons that include *Z. atricapilla.* High *F*_ST_ windows are associated with low to moderate *π*_Between_ (Fig. 4B). There is also a much stronger correlation between *π*_Between_ and *π*_Within_ in this comparison (Fig. 4C) than in the comparisons involving *Z. atricapilla* (Figs. 2C and 3C). These patterns are consistent with a history of selection in the ancestral population followed by divergent selection at these loci in daughter populations, or introgression of adaptive alleles between populations, followed by divergent selection on those loci (Cruickshank & Hahn, 2014; Irwin et al., 2016, 2018).

In contrast to the comparisons involving *Z. atricapilla*, which showed broad regions of increased differentiation, differentiation peaks between *Z. l. gambelii* and *Z. l. pugetensis* are short and localized, aside from one large block on chromosome 1A (discussed below). The whole-genome and per chromosome PCAs show clear evidence of hybridization between *Z. l. gambelii* and *Z. l. pugetensis*, with one back-crossed individual in the dataset, and several others that show evidence of introgression. Considering the limited sample sizes in this study, this suggests that hybridization is more common between these subspecies than one might expect from their allopatric distributions. A block on chromosome 1A shows the highest autosomal *F*_ST_ between *Z. l. gambelii* and *Z. l. pugetensis*. Furthermore, three individuals are heterozygous for different haplotypes across this block despite being otherwise genetically *Z. l. pugetensis*, which suggests that recombination is suppressed in this region. The increase in LD between the groups relative to within groups in this region is only slight, suggesting that if recombination in the region is suppressed due to an inversion, it would have occurred very recently, as the inversion haplotypes would not have had sufficient time to differentiate and develop strong LD between loci. Across this block, *π*_Within_ is slightly reduced in both species and is concordant with an overall level of *π*_Between_ that appears similar to the rest of the genome (Fig S5), suggesting that differentiation across this region could also be the result of divergent selection in an area of reduced recombination. Chromosome 1A has been implicated in divergence in other avian systems (Baiz et al., 2020; Irwin et al., 2018; Wang et al., 2020) and warrants further investigation in this taxon.

In all comparisons, autosomal differentiation tends to be higher on larger autosomes (Fig 5). Relative differentiation increases with chromosome length, and larger autosomes tend to have broader islands of differentiation while smaller chromosomes generally show narrow peaks of differentiation, if any at all (Figs 2A, 3A, and 4A). This is consistent with patterns observed in the Chicken (*Gallus gallus*) genome (Mugal et al., 2013) and the White-throated Sparrow (*Zonotrichia albicolis*) genome (Huynh et al., 2010), which is closely related to the *Zonotrichia* sparrows investigated here. The avian microchromosomes experience a higher recombination rate than the macrochromosomes, and are thus predicted to have higher diversity (Groenen et al., 2009; Mugal et al., 2013). This may also explain why peaks of differentiation on the microchromosomes are narrower than those on larger chromosomes, as higher recombination rates would reduce the strength of genetic hitchhiking.

Comparisons of *F*_ST_, *π*_Between_, and *π*_Within_ across the Z chromosome and autosomes suggest that the Z chromosome has played an outsized role in divergence between *Z. atricapilla* and *Z. leucophrys*. While increased relative differentiation on the Z chromosome relative to autosomes has been observed in many avian species pairs, the *F*_ST,Z_ / *F*_ST, A_ ratios for comparisons involving *Z. atricapilla* are high. In contrast, the ratio for the two *Z*. *leucophrys* is similar to those found in other passerines (Irwin, 2018). Increased *F*_ST_ across the Z chromosome is typically driven by decreased *π*_Within_ since the effective population size of the Z chromosome tends to be reduced relative to the autosomes (Charlesworth, 2001; Ellegren, 2009). Ratios of *π*_Within, Z*_ / *π*_Within, A_ that fall below the value of 0.5625 cannot be explained merely by a neutral model with extreme variance in male reproductive success (Charlesworth, 2001; Irwin, 2018). The ratios of *π*_Within, Z*_ / *π*_Within, A_ were below 1 in all taxa in this study and significantly below 0.5625 for both *Z. atricapilla* and *Z. l. leucophrys*, suggesting increased impact of diversity-reducing selection on the Z chromosome relative to the autosomes in both these taxa. It should be noted that a ratio of *π*_Within, Z*_ / *π*_Within, A_ of 0.5625 only occurs without selection when there is extreme variance in reproductive success of males, such as in lekking species. While some studies have found that upwards of 38% of *Z. leucophrys* nestlings were the result of extra-pair paternity (Sherman & Morton, 1988), the majority are monogamous breeders with a high degree of parental care provided by the male (Norment, 1992). Hence, while the observed *π*_Within, Z*_/ *π*_Within, A_ ratio for Z. *l. gambelii* of 0.5469 was not significantly below 0.5625, we still view this as evidence that the Z chromosome may have been subject to selection in this lineage as well. Regardless of the evolutionary force driving it, reduced *π*_Within, Z*_ contributes to the increased differentiation of the Z chromosome in all three groups. Furthermore, the extreme increase in *F*_ST_ in the comparisons involving *Z. atricapilla* is also driven by an increase in *π*_Between, Z_ that is not observed in the comparison between the two *Z. leucophrys* groups. In other species pairs, *π*_Between, Z_ is often reduced relative to *π*_Between, A_ (Irwin et al., 2016, 2018), possibly due to reduced effective population size of the Z chromosome in the common ancestor. While this is the case when comparing *Z. l. gambelii* and *Z. l. pugetensis, π*_Between, Z_ was increased relative to the autosomes in comparisons between *Z. atricapilla* and *Z. leucophrys.* This suggests reduced gene flow, and an increased coalescence time between these species for the Z chromosome compared to the autosomes (Cruickshank & Hahn, 2014).

These patterns are consistent with a large chromosomal inversion spanning most of the Z chromosome, with inversion haplotypes segregating between *Z. atricapilla* and *Z. leucophrys*. Increased *π*_Between, Z_ suggests decreased gene flow, which could be explained by a large chromosomal inversion repressing recombination between the Z chromosomes for an extended period of time, even when there was still substantial gene flow between these species at other parts of the genome. *F*_ST_ increases sharply and remains elevated across most of the chromosome, and the genotypes of individuals across this region are consistently hemi or homozygous for a species-specific haplotype. Furthermore, linkage disequilibrium plots suggest that recombination is suppressed across a broad area of the Z chromosome between the two species, but not within either species. This pattern in linkage disequilibrium is similar to those observed in other systems across chromosomal inversions (Hooper et al., 2019; Todesco et al., 2020a). Despite the high level of synteny of the avian genome, rearrangements and inversions within the Z chromosome are common (Backström et al., 2006; Yazdi & Ellegren, 2018), and Z-linked inversions have a higher fixation rate than autosomal inversions (Hooper & Price, 2017). Thus, an inversion is a plausible explanation for the observed patterns. The regions of reduced *F*_ST_ and LD within the putative inversion may represent translocations relative to the reference genome, given the long history of independent evolution between *Zonotrichia* and the Zebra Finch (Oliveros et al., 2019).

Inversions are challenging to investigate with the short-read and reduced representation sequencing techniques used in this study. Further evidence for an inversion could be provided by aligning reference genomes constructed from these two species to one another (e.g. Todesco et al., 2020b), but constructing such a reference genome is beyond the scope of this study. Evidence could also be provided by analyzing heterozygosity and linkage disequilibrium across the putative inversion in hybrids (Hooper et al., 2019). This may be a challenge for the putative Z chromosome inversion, as hybrids are rare, and there is no specific region in which hybridization is more likely to occur. The Iona Island Bird Observatory in Richmond, British Columbia, was selected as the primary sampling site for this study based on capturing two putative *Z. leucophrys × atricapilla* hybrids during the previous spring migration monitoring period, when genetic samples were not being collected. Consistent with previous descriptions of putative hybrids (Miller, 1940), these individuals showed intermediate crown phenotype between the species, with a medial yellow crown stripe similar to *Z. atricapilla* and the white supercilium of *Z. leucophrys* (Fig S8). Unfortunately, we did not capture any hybrids during our field season for the present study, but IIBO is located along the pacific flyway, a common migratory route for songbirds, and regularly captures both *Z. atricapilla* and *Z. l. gambelii* on migration (Lisovski et al., 2019), as well as *Z. l. pugetensis* which can be found in the Lower Mainland year-round (Campbell et al., 2001). Since the sympatric zone between *Z. atricapilla* and *Z. leucophrys* is vast, migration stopover sites like IIBO may be ideal places to sample hybrids, as many individuals concentrate in these sites during migration. Genomic analysis of these putative hybrids could further elucidate the nature of reproductive isolation between *Z. atricapilla* and *Z. leucophrys*. For example, if all putative hybrids are F1s, it would suggest that they are not able to reproduce, especially given the lack of evidence for current gene flow in our genomic analysis of these two species.

Given the evidence of a chromosomal inversion on the Z chromosome, what is the possible significance of this structural variant to the history and future of speciation in this system? Patterns of differentiation across the autosomes as well as the distributions of these species (Weir & Schluter, 2004) are consistent with periods of geographic separation and differentiation, likely during past glacial maxima. In one of the two incipient species during this period, the inversion may have arisen and become fixed. Upon secondary contact, the inversion would suppress recombination and prevent local introgression between the two inversion haplotypes (Rieseberg, 2001). Any genes within the inversion that contribute to isolation in these species would be tightly linked and essentially act additively as a single locus (Rieseberg, 2001). This lack of recombination and introgression would also allow for further differentiation and accumulation of genetic incompatibilities that further reduce the hybrid fitness (Noor et al., 2001). Given the number of genes on the Z chromosome implicated in plumage pigmentation (e.g. Campagna et al., 2017; Sæther et al., 2007; Toews, Taylor, et al., 2016), a key component of avian sexual signalling and species recognition, it is possible that this inversion has captured differential alleles for some of these genes, and thus may be involved in pre-mating isolation between the two species. Large inversions such as this are also likely to capture loci involved in genetic incompatibilities. Such incompatibilities are relatively common (Noor et al., 2001; Presgraves, 2003), which would contribute to post-mating isolation between these species. Thus, the putative inversion reported here may contribute to both pre-and post-mating isolation, and may have helped prevent the homogenization of the genome and collapse of these species during past secondary contact.

This study found that *Z. atricapilla* and *Z. leucophrys* show high nuclear genomic differentiation that varies greatly among different parts of the genome, pointing to a long history of divergence due in part to geographic separation but with some periods of gene flow. The regions of high nuclear differentiation contrast markedly with the close similarity in mitochondrial DNA, pointing to somewhat recent hybridization, introgression, and replacement of mitochondrial DNA (Taylor et al., 2021). Despite this evidence for mitochondrial gene flow and their current broadly overlapping ranges, we found no evidence of current nuclear gene flow between *Z. atricapilla* and *Z. leucophrys*, suggesting a recent cut-off of gene flow between the species. In contrast, the subspecies *Z. l. gambelii* and *Z. l. pugetensis* show lower genomic differentiation, patterns of nucleotide diversity characteristic of recurrent selection in ancestral and daughter populations, and evidence of ongoing gene flow between the two groups. Differentiation between all three taxa is particularly strong on the Z chromosome, and distinct Z chromosome inversion haplotypes that segregate between *Z. atricapilla* and *Z. leucophrys* may have driven early divergence between these groups and contributed to reproductive isolation between them. This finding adds to the growing body of evidence for the outsized role of inversions in divergence between taxa, and further investigation into this species pair could allow us to better understand the role of inversions in speciation.

## Acknowledgments

We would like to acknowledge and thank WildResearch, especially the staff and volunteers at the Iona Island Bird Observatory, as well as Madelyn Ore, Ben Freeman, Julian Heavyside, Jen Muñoz, Kimberly Dohms, Matthias Bieber, Max Edworthy, Daniel Froehlich, and Coni Rivas, for their help with field sample collection. Thanks to Ildiko Szabo for additional samples from the Cowan Tetrapod Collection. Thank you to the rest of the Irwin Lab group for the feedback and thought-provoking discussions. Thank you to Metro Vancouver Parks for permission to collect samples from IIBO, and the Federal Bird Banding Office at Environment and Climate Change Canada for the permit to take blood samples (permit number 10746 P).

## Author Contributions

KA, QM, and DI conceived the study; AH coordinated fieldwork with IIBO; QM, FF, and LN collected the samples; QM, FF, EN, and LN performed DNA extractions and constructed GBS libraries; QM analyzed the data with advice from KA and DI; QM, KA and DI wrote the manuscript; all authors contributed to drafts and gave final publication approval.

## Literature Cited

Alcaide, M., Scordato, E. S. C., Price, T. D., & Irwin, D. E. (2014). Genomic divergence in a ring species complex. Nature, 511(7507), 83–85. https://doi.org/10.1038/nature13285

Backström, N., Brandström, M., Gustafsson, L., Qvarnström, A., Cheng, H., & Ellegren, H. (2006). Genetic mapping in a natural population of collared flycatchers (Ficedula albicollis): Conserved synteny but gene order rearrangements on the avian Z chromosome. Genetics, 174(1), 377–386. https://doi.org/10.1534/genetics.106.058917

Baiz, M. D., Wood, A. W., Brelsford, A., Lovette, I. J., & Toews, D. P. L. (2020). Pigmentation Genes Show Evidence of Repeated Divergence and Multiple Bouts of Introgression in Setophaga Warblers. Current Biology, 1–7. https://doi.org/10.1016/j.cub.2020.10.094

Ballard, J. W., & Whitlock, M. C. (2004). The incomplete natural history of mitochondria. Molecular Ecology, 13, 729–744.

Baute, G. J., Owens, G. L., Bock, D. G., & Rieseberg, L. H. (2016). Genome-wide genotyping-by-sequencing data provide a high-resolution view of wild helianthus diversity, genetic structure, and interspecies gene flow. American Journal of Botany, 103(12), 2170–2177. https://doi.org/10.3732/ajb.1600295

Bolger, A. M., Lohse, M., & Usadel, B. (2014). Trimmomatic: a flexible trimmer for Illumina sequence data. Bioinformatics, 30(15), 2114–2120. https://doi.org/10.1093/bioinformatics/btu170

Campagna, L., Repenning, M., Silveira, L. F., Fontana, C. S., Tubaro, P. L., & Lovette, I. J. (2017). Repeated divergent selection on pigmentation genes in a rapid finch radiation. Science Advances, 3(5), 1–12. https://doi.org/10.1126/sciadv.1602404

Chang, C. C., Chow, C. C., Tellier, L. C. A. M., Vattikuti, S., Purcell, S. M., & Lee, J. J. (2015). Second-generation PLINK: Rising to the challenge of larger and richer datasets. GigaScience, 4(1), 1–16. https://doi.org/10.1186/s13742-015-0047-8

Charlesworth, B. (2001). The effect of life-history and mode of inheritance on neutral genetic variability. Genetical Research, 77(2), 153–166. https://doi.org/10.1017/S0016672301004979

Chilton, G., Baker, M. C., Barrentine, C. D., & Cunningham, M. A. (2020). White-crowned Sparrow (Zonotrichia leucophrys), version 1.0. In Birds of the World (A. F. Poole and F. B. Gill, Editors). Cornell Lab of Ornithology. https://doi.org/10.2173/bow.whcspa.01

Cowles, S. A., & Uy, J. A. C. (2019). Rapid, complete reproductive isolation in two closely related Zosterops White-eye bird species despite broadly overlapping ranges*. Evolution, 73(8), 1647–1662. https://doi.org/10.1111/evo.13797

Cruickshank, T. E., & Hahn, M. W. (2014). Reanalysis suggests that genomic islands of speciation are due to reduced diversity, not reduced gene flow. Molecular Ecology, 23(13), 3133–3157. https://doi.org/10.1111/mec.12796

Danecek, P., Auton, A., Abecasis, G., Albers, C. A., Banks, E., DePristo, M. A., Handsaker, R. E., Lunter, G., Marth, G. T., Sherry, S. T., McVean, G., & Durbin, R. (2011). The variant call format and VCFtools. Bioinformatics, 27(15), 2156–2158. https://doi.org/10.1093/bioinformatics/btr330

Ellegren, H. (2009). The different levels of genetic diversity in sex chromosomes and autosomes. Trends in Genetics, 25(6), 278–284. https://doi.org/10.1016/j.tig.2009.04.005

Elshire, R. J., Glaubitz, J. C., Sun, Q., Poland, J. A., Kawamoto, K., Buckler, E. S., & Mitchell, S. E. (2011). A robust, simple genotyping-by-sequencing (GBS) approach for high diversity species. PloS ONE, 6(5), 1–10. https://doi.org/10.1371/journal.pone.0019379

Geraldes, A., Askelson, K. K., Nikelski, E., Doyle, F. I., Harrower, W. L., Winker, K., & Irwin, D. E. (2019). Population genomic analyses reveal a highly differentiated and endangered genetic cluster of northern goshawks (Accipiter gentilis laingi) in Haida Gwaii. Evolutionary Applications, 12(4), 757–772. https://doi.org/10.1111/eva.12754

Groenen, M. A. M., Wahlberg, P., Foglio, M., Cheng, H. H., Megens, H. J., Crooijmans, R. P. M. A., Besnier, F., Lathrop, M., Muir, W. M., Wong, G. K. S., Gut, I., & Andersson, L. (2009). A high-density SNP-based linkage map of the chicken genome reveals sequence features correlated with recombination rate. Genome Research, 19(3), 510–519. https://doi.org/10.1101/gr.086538.108

Grossen, C., Seneviratne, S. S., Croll, D., & Irwin, D. E. (2016). Strong reproductive isolation and narrow genomic tracts of differentiation among three woodpecker species in secondary contact. Molecular Ecology, 25(17), 4247–4266. https://doi.org/10.1111/mec.13751

Hill, G. E. (2019). Mitonuclear Ecology. Oxford University Press.

Hooper, D. M., Griffith, S. C., & Price, T. D. (2019). Sex chromosome inversions enforce reproductive isolation across an avian hybrid zone. Molecular Ecology, 28(6), 1246–1262. https://doi.org/10.1111/mec.14874

Hooper, D. M., & Price, T. D. (2017). Chromosomal inversion differences correlate with range overlap in passerine birds. Nature Ecology and Evolution, 1(10), 1526–1534. https://doi.org/10.1038/s41559-017-0284-6

Hunn, E. S., & Beaudette, D. (2014). Apparent sympatry of two subspecies of the White-Crowned Sparrow, Zonotrichia leucophrys pugetensis and gambelii, in Washington state. Western Birds, 45, 132–140.

Huynh, L. Y., Maney, D. L., & Thomas, J. W. (2010). Contrasting population genetic patterns within the white-throated sparrow genome (Zonotrichia albicollis). BMC Genetics, 11(1), 96. https://doi.org/10.1186/1471-2156-11-96

Irwin, D. E. (2018). Sex chromosomes and speciation in birds and other ZW systems. Molecular Ecology, 27(19), 3831–3851. https://doi.org/10.1111/mec.14537

Irwin, D. E., Rubtsov, A. N., & Panov, E. N. (2009). Mitochondrial introgression and replacement between yellowhammers (Emberiza 18itronella) and pine buntings (Emberiza leucocephalos) (Aves: Passeriformes). Biological Journal of the Linnean Society, 98, 422–438.

Irwin, D. E., Alcaide, M., Delmore, K. E., Irwin, J. H., & Owens, G. L. (2016). Recurrent selection explains parallel evolution of genomic regions of high relative but low absolute differentiation in a ring species. Molecular Ecology, 25(18), 4488–4507. https://doi.org/10.1111/mec.13792

Irwin, D. E., Milá, B., Toews, D. P. L., Brelsford, A., Kenyon, H. L., Porter, A. N., Grossen, C., Delmore, K. E., Alcaide, M., & Irwin, J. H. (2018). A comparison of genomic islands of differentiation across three young avian species pairs. Molecular Ecology, 27(23), 4839–4855. https://doi.org/10.1111/mec.14858

Li, H., & Durbin, R. (2009). Fast and accurate short read alignment with Burrows-Wheeler transform. Bioinformatics, 25(14), 1754–1760. https://doi.org/10.1093/bioinformatics/btp324

Li, H., Handsaker, B., Wysoker, A., Fennell, T., Ruan, J., Homer, N., Marth, G., Abecasis, G., & Durbin, R. (2009). The Sequence Alignment/Map format and SAMtools. Bioinformatics, 25(16), 2078–2079. https://doi.org/10.1093/bioinformatics/btp352

Lowry, D. B., & Willis, J. H. (2010). A widespread chromosomal inversion polymorphism contributes to a major life-history transition, local adaptation, and reproductive isolation. PloS Biology, 8(9). https://doi.org/10.1371/journal.pbio.1000500

Mank, J. E., Vicoso, B., Berlin, S., & Charlesworth, B. (2010). Effective population size and the Faster-X effect: empirical results and their interpretation. Evolution, 64, 663–674.

McKenna, A., Hanna, M., Banks, E., Sivachenko, A., Cibulskis, K., Kernytsky, A., Garimella, K., Altshuler, D., Gabrial, S., Daly, M., & DePristo, M. A. (2010). The Genome Analysis Toolkit: A MapReduce framework for analyzing next-generation DNA sequencing data. Genome Research, 20, 1297–1303. https://doi.org/10.1101/gr.107524.110.

Mikkelsen, E. K., & Irwin, D. E. (2021). Ongoing production of low-fitness hybrids limits range overlap between divergent cryptic species. BioRxiv Preprint, 2008, 1–13.

Miller, A. H. (1940). A Hybrid between Zonotrichia coronata and Zonotrichia leucophrys. The Codor, 42(1), 45–48.

Morton, M. L., & Mewaldt, L. R. (1960). Further Evidence of Hybridization between Zonotrichia atricapilla and Zonotrichia leucophrys. The Condor: Ornithological Applications, 62(6), 485–486. https://doi.org/10.1093/condor/62.6.485

Mugal, C. F., Nabholz, B., & Ellegren, H. (2013). Genome-wide analysis in chicken reveals that local levels of genetic diversity are mainly governed by the rate of recombination. BMC Genomics, 14(1). https://doi.org/10.1186/1471-2164-14-86

Navarro, A., & Barton, N. H. (2003). Accumulating postzygotic isolation genes in parapatry: A new twist on chromosomal speciation. Evolution, 57(3), 447–459. https://doi.org/10.1111/j.0014-3820.2003.tb01537.x

Noor, M. A. F., Gratos, K. L., Bertucci, L. A., & Reiland, J. (2001). Chromosomal inversions and the reproductive isolation of species. Proceedings of the National Academy of Sciences of the United States of America, 98(21), 12084–12088. https://doi.org/10.1073/pnas.221274498

Norment, C. J., Hendricks, P., & Santonocito, R. (2020). Golden-crowned Sparrow (Zonotrichia atricapilla), version 1.0. In Birds of the World (A. F. Poole and F. B. Gill, Editors). Cornell Lab of Ornithology. https://doi.org/10.2173/bow.gocspa.01

Oliveros, C. H., Field, D. J., Ksepka, D. T., Keith Barker, F., Aleixo, A., Andersen, M. J., Alström, P., Benz, B. W., Braun, E. L., Braun, M. J., Bravo, G. A., Brumfield, R. T., Terry Chesser, R., Claramunt, S., Cracraft, J., Cuervo, A. M., Derryberry, E. P., Glenn, T. C., Harvey, M. G.,…Faircloth, B. C. (2019). Earth history and the passerine superradiation. Proceedings of the National Academy of Sciences of the United States of America, 116(16), 7916–7925. https://doi.org/10.1073/pnas.1813206116

Ortiz-Barrientos, D., Engelstädter, J., & Rieseberg, L. H. (2016). Recombination Rate Evolution and the Origin of Species. Trends in Ecology and Evolution, 31(3), 226–236. https://doi.org/10.1016/j.tree.2015.12.016

Owens, G. L., Baute, G. J., & Rieseberg, L. H. (2016). Revisiting a classic case of introgression: hybridization and gene flow in Californian sunflowers. Molecular Ecology, 25(11), 2630–2643. https://doi.org/10.1111/mec.13569

Presgraves, D. C. (2003). A fine-scale genetic analysis of hybrid incompatibilities in Drosophila. Genetics, 163(3), 955–972. https://doi.org/10.1093/genetics/163.3.955

Presgraves, D. C. (2018). Evaluating genomic signatures of “the large X-effect” during complex speciation. Molecular Ecology, 27, 3822–3830.

Rhie, A., McCarthy, S. A., Fedrigo, O., Damas, J., Formenti, G., Koren, S., Uliano-Silva, M., Chow, W., Fungtammasan, A., Kim, J., Lee, C., Ko, B. J., Chaisson, M., Gedman, G. L., Cantin, L. J., Thibaud-Nissen, F., Haggerty, L., Bista, I., Smith, M.,…Jarvis, E. D. (2021). Towards complete and error-free genome assemblies of all vertebrate species. Nature, 592(7856), 737–746. https://doi.org/10.1038/s41586-021-03451-0

Rieseberg, L. H. (2001). Chromosomal rearrangements and speciation. TRENDS in Ecology & Evolution, 16(7), 351–358. http://tree.trends.com

Sæther, S. A., Sætre, G.-P., Borge, T., Wiley, C., Svedin, N., Andersson, G., Veen, T., Haavie, J., Servedio, M. R., Bureš, S., Král, M., Hjernquist, M. B., Gustafsson, L., Träff, J., & Qvarnström, A. (2007). Sex chromosome-linked species recognition and evolution of reproductive isolation in flycatchers. Science, 318(5847), 95–97.

Sherman, P. W., & Morton, M. L. (1988). Extra-pair fertilizations in mountain white-crowned sparrows. Behavioral Ecology and Sociobiology, 22(6), 413–420. https://doi.org/10.1007/BF00294979

Seutin, G., White, B. N., & Boag, P. T. (1991). Preservation of avian blood and tissue samples for DNA analyses. Canadian Journal of Zoology, 69(1), 82–90. https://doi.org/10.1139/z91-013

Stacklies W, Redestig H, Scholz M, Walther D, Selbig J (2007) PcaMethods – a Bioconductor package providing PCA methods for incomplete data. Bioinformatics 23:1164–1167

Taylor, R. S., Bramwell, A. C., Clemente-Carvalho, R., Cairns, N. A., Bonier, F., Dares, K., & Lougheed, S. C. (2021). Cytonuclear discordance in the crowned-sparrows, Zonotrichia atricapilla and Zonotrichia leucophrys. Molecular Phylogenetics and Evolution, 162, 107216. https://doi.org/10.1016/j.ympev.2021.107216

Cheung, W., Staton, S. E., Muños, S., Nielsen, R., Donovan, L. A.,…Rieseberg, L. H. (2020b). Massive haplotypes underlie ecotypic differentiation in sunflowers. Nature, 584(7822), 602–607. https://doi.org/10.1038/s41586-020-2467-6

Toews, D. P. L., & Brelsford, A. (2012). The biogeography of mitochondrial and nuclear discordance in animals. Molecular Ecology, 21(16), 3907–3930. https://doi.org/10.1111/j.1365-294X.2012.05664.x

Toews, D. P. L., Campagna, L., Taylor, S. A., Balakrishnan, C. N., Baldassarre, D. T., Deane-Coe, P. E., Harvey, M. G., Hooper, D. M., Irwin, D. E., Judy, C. D., Mason, N. A., McCormack, J. E., McCracken, K. G., Oliveros, C. H., Safran, R. J., Scordato, E. S. C., Stryjewski, K. F., Tigano, A., Uy, J. A. C., & Winger, B. M. (2016). Genomic approaches to understanding population divergence and speciation in birds. The Auk, 133(1), 13–30. https://doi.org/10.1642/auk-15-51.1

Toews, D. P. L., Taylor, S. A., Vallender, R., Brelsford, A., Butcher, B. G., Messer, P. W., & Lovette, I. J. (2016). Plumage Genes and Little Else Distinguish the Genomes of Hybridizing Warblers. Current Biology, 26(17), 2313–2318. https://doi.org/10.1016/j.cub.2016.06.034

Thomas, J. W., Cáceres, M., Lowman, J. J., Morehouse, C. B., Short, M. E., Baldwin, E. L., Maney, D. L., & Martin, C. L. (2008). The chromosomal polymorphism linked to variation in social behavior in the white-throated sparrow (Zonotrichia albicollis) is a complex rearrangement and suppressor of recombination. Genetics, 179(3), 1455–1468. https://doi.org/10.1534/genetics.108.088229

Todesco, M., Owens, G. L., Bercovich, N., Légaré, J. S., Soudi, S., Burge, D. O., Huang, K., Ostevik, K. L., Drummond, E. B. M., Imerovski, I., Lande, K., Pascual-Robles, M. A., Nanavati, M., Jahani, M., Cheung, W., Staton, S. E., Muños, S., Nielsen, R., Donovan, L. A.,…Rieseberg, L. H. (2020a). Massive haplotypes underlie ecotypic differentiation in sunflowers. Nature, 584(7822), 602–607. https://doi.org/10.1038/s41586-020-2467-6

Todesco, M., Owens, G. L., Bercovich, N., Légaré, J. S., Soudi, S., Burge, D. O., Huang, K., Ostevik, K. L., Drummond, E. B. M., Imerovski, I., Lande, K., Pascual-Robles, M. A., Nanavati, M., Jahani, M., Wang, S., Ore, M. J., Mikkelsen, E. K., Lee-Yaw, J., Toews, D. P. L., Rohwer, S., & Irwin, D. (2021). Signatures of mitonuclear coevolution in a warbler species complex. Nature Communications, 12: 4279.

Wang, S., Rohwer, S., Zwaan, D. R. De, Toews, D. P. L., Lovette, I. J., Mackenzie, J., & Irwin, D. E. (2020). Selection on a small genomic region underpins differentiation in multiple color traits between two warbler species. Evolution Letters, 1–14. https://doi.org/10.1002/evl3.198

Warren, W. C., Clayton, D. F., Ellegren, H., Arnold, A. P., Hillier, L. W., Künstner, A., Searle, S., White, S., Vilella, A. J., Fairley, S., Heger, A., Kong, L., Ponting, C. P., Jarvis, E. D., Mello, C. V., Minx, P., Lovell, P., Velho, T. A. F., Ferris, M.,…Wilson, R. K. (2010). The genome of a songbird. Nature, 464(7289), 757–762. https://doi.org/10.1038/nature08819

Weckstein, J. D., Zink, R. M., Blackwell-Rago, R. C., & Nelson, D. A. (2001). Anomalous Variation in Mitochondrial Genomes of White-Crowned (Zonotrichia leucophrys) and Golden-Crowned (Z. atricapilla) Sparrows: Pseudogenes, Hybridization, or Incomplete Lineage Sorting? The Auk, 118(1), 231–236. https://doi.org/10.2307/4089773

Weir, B. S., & Cockerham, C. C. (1984). Estimating F-statistics for the analysis of population structure. Evolution, 38(6), 1358–1370. https://doi.org/10.1111/j.1558-5646.1984.tb05657.x

Weir, J. T., & Schluter, D. (2004). Ice sheets promote speciation in boreal birds. Proceedings of the Royal Society B: Biological Sciences, 271(1551), 1881–1887. https://doi.org/10.1098/rspb.2004.2803

Yazdi, H. P., & Ellegren, H. (2018). A genetic map of ostrich Z chromosome and the role of inversions in avian sex chromosome evolution. Genome Biology and Evolution, 10(8), 2049–2060. https://doi.org/10.1093/gbe/evy163

Zink, R. M., & Blackwell, R. C. (1996). Patterns of allozyme, mitochondrial DNA, and morphometric variation in four sparrow genera. The Auk, 113(1), 59–67.

Zink, R. M., Dittmann, D. L., & Rootes, W. L. (1991). Mitochondrial DNA Variation and the Phylogeny of Zonotrichia. The Auk, 108(3), 578–584.

