## Supplementary Figures for "Extreme sex chromosome differentiation, likely driven by inversion, contrasts with mitochondrial paraphyly between species of crowned sparrows"

### Supplementary Material

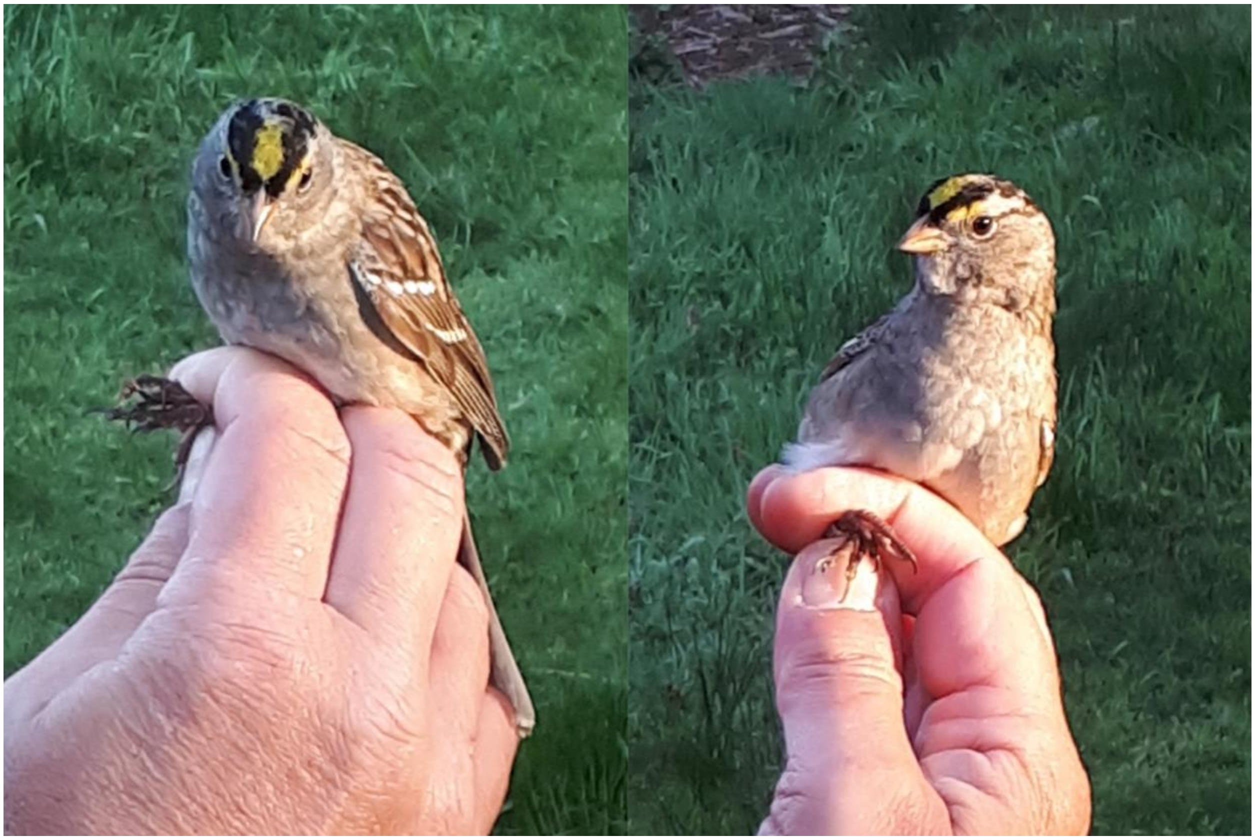

**Figure S1.** A putative *Zonotrichia atricapilla* x *leucophrys* hybrid captured at the Iona Island Bird Observatory on May 2, 2018, during migration monitoring. Another individual (not shown) was captured on May 8, 2018.

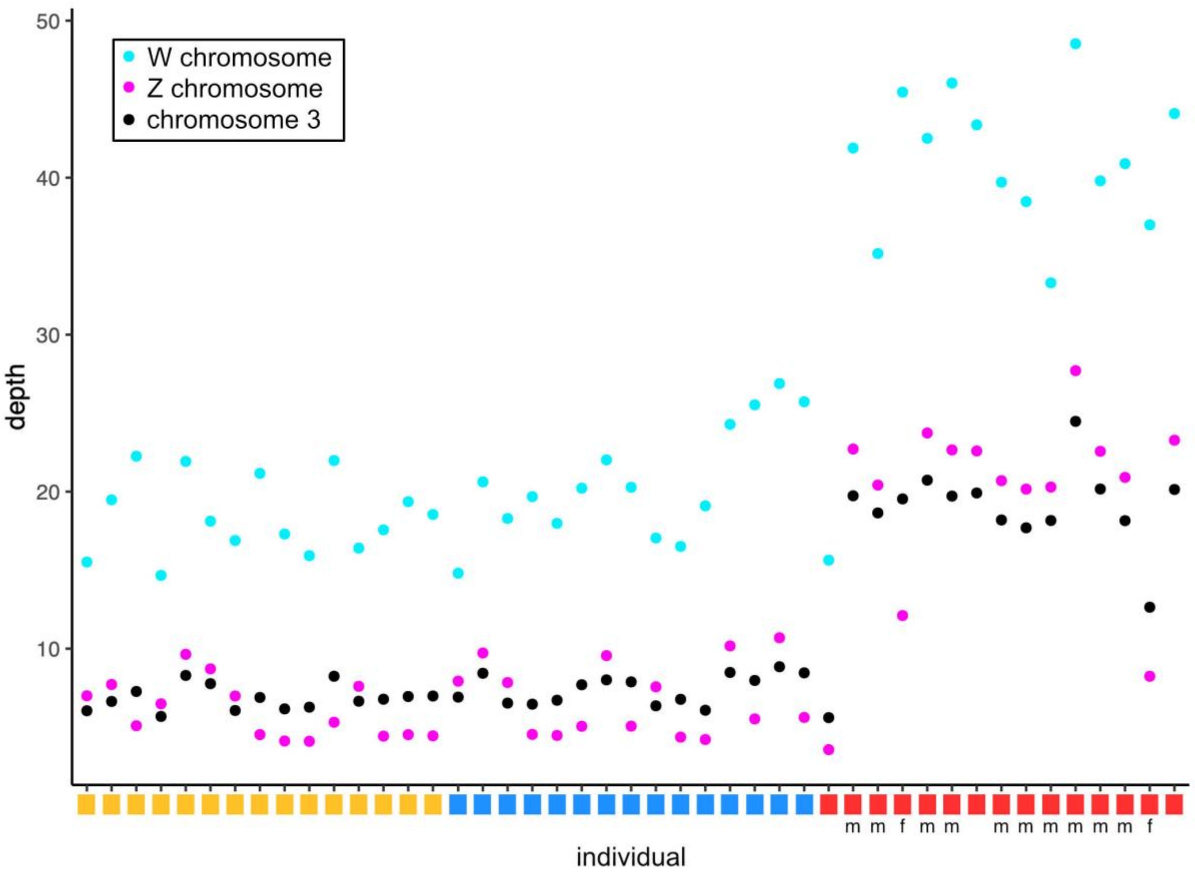

**Figure S2.** Reads mapping to the Zebra Finch W chromosome show anomalously high depth of coverage for all individuals**.** Mean depth of coverage across reads mapping to chromosome 3 (black), the Z chromosome (magenta), and the W chromosome (cyan) in the Zebra Finch reference genome for each individual. The taxon of each individual is indicated by the coloured rectangle, with yellow representing *Z. atricapilla*, blue representing *Z. l. gambelii*, and red representing *Z. l. pugetensis.* For individuals where sex was inferred from breeding characteristics during sampling, m indicates male and f indicates female. Note that most *Z. l. pugetensis* samples were sequenced on an Illumina NovaSeq 6000, while the remainder of the samples were sequenced on an Illumina HiSeq 4000, which accounts for the higher read depth for these individuals.

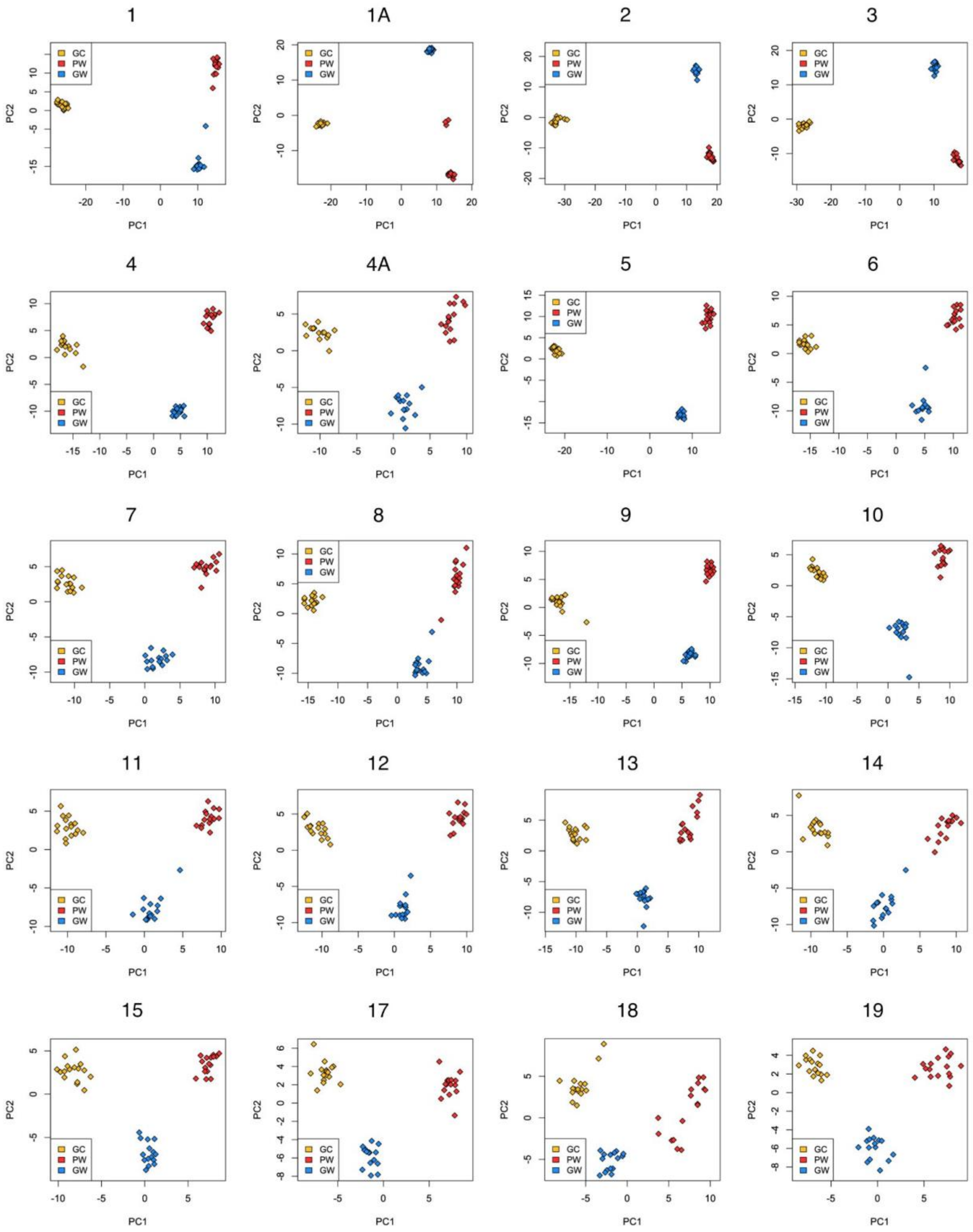
1
1A
2
3
4
4A
17
18
19
GC
PW
GW
PC1
PC2

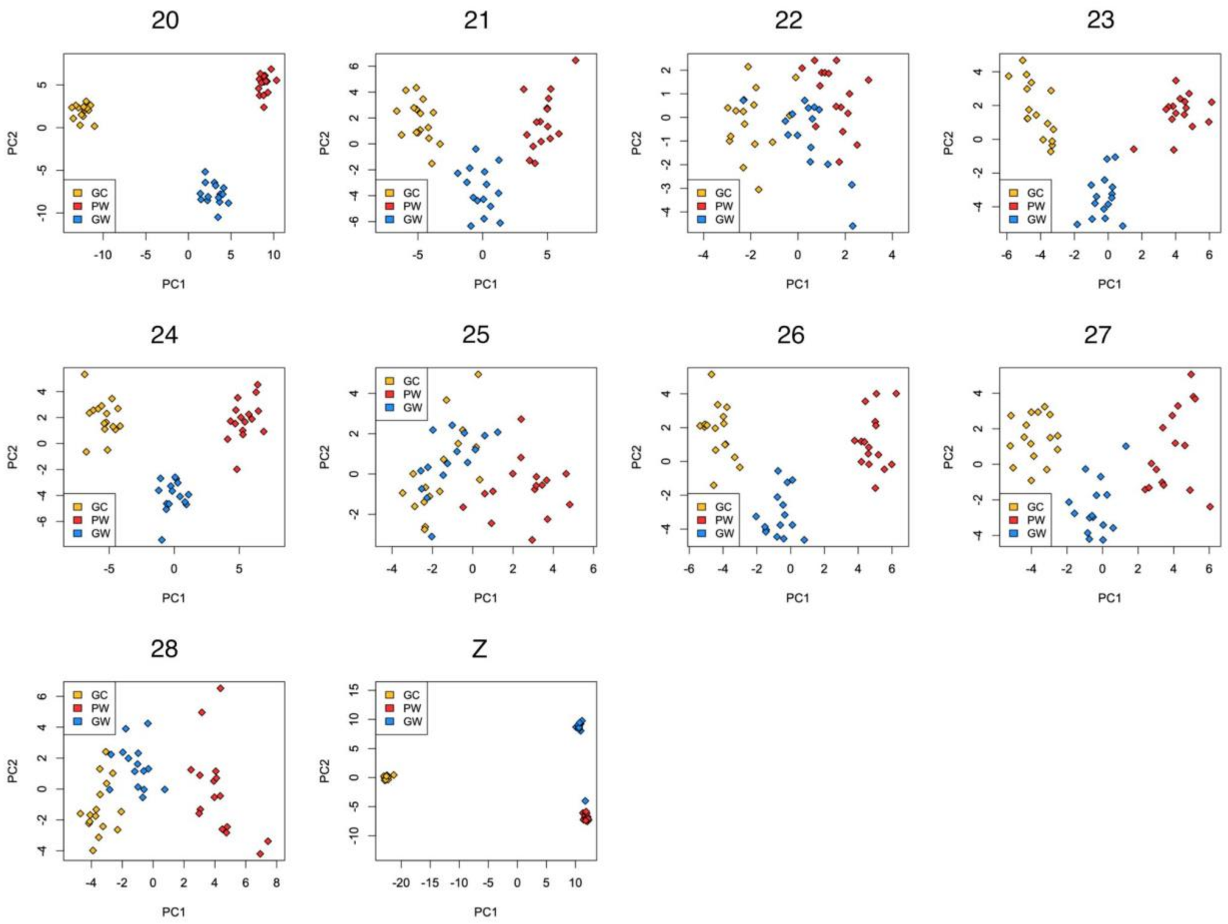

**Figure S3.** Principal Component Analyses of all samples for SNPs on each chromosome. Yellow represents *Z. atricapilla* individuals, blue represents *Z. l. gambelii*, and red represents *Z. l. pugetensis*.

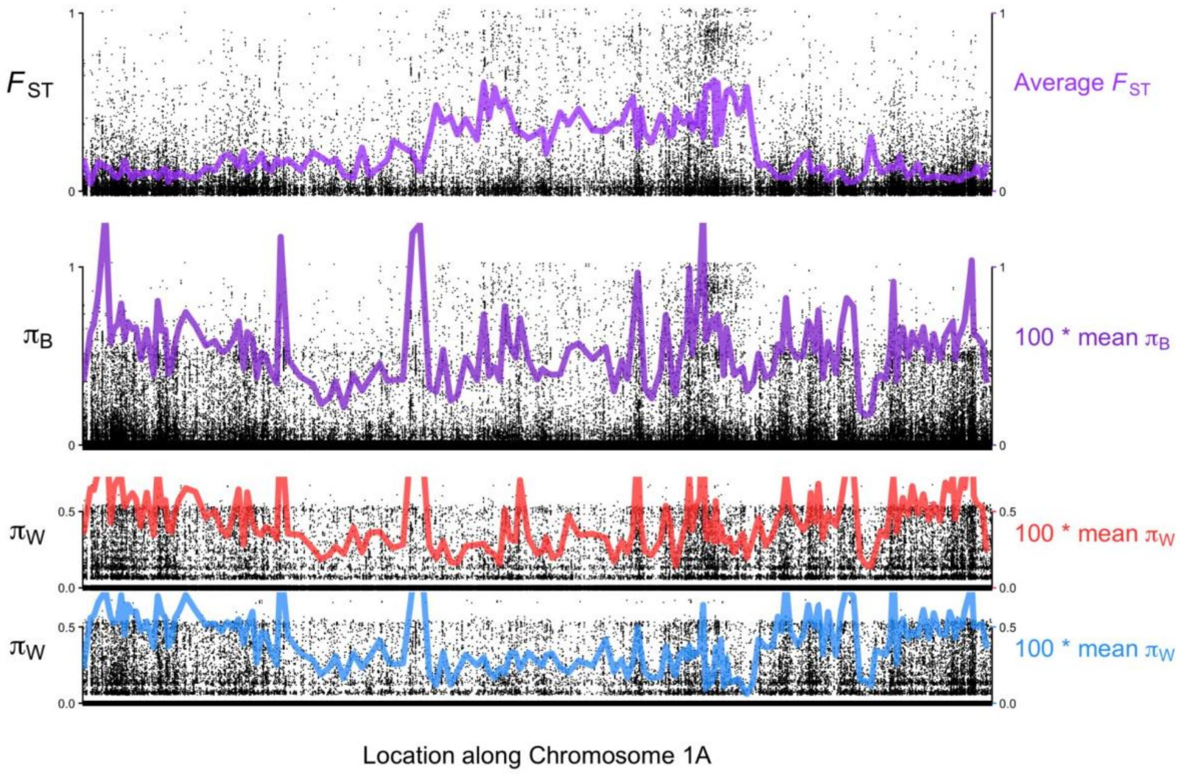

**Figure S4.** Patterns of $F_{ST}$, $\pi_{Between}$, and $\pi_{Within}$ across chromosome 1A in comparisons between *Z. l. gambelii* (blue) and *Z. l. pugetensis* (red). Black dots represent individual SNPs, while coloured lines represent average across 10,000 bp sliding windows.

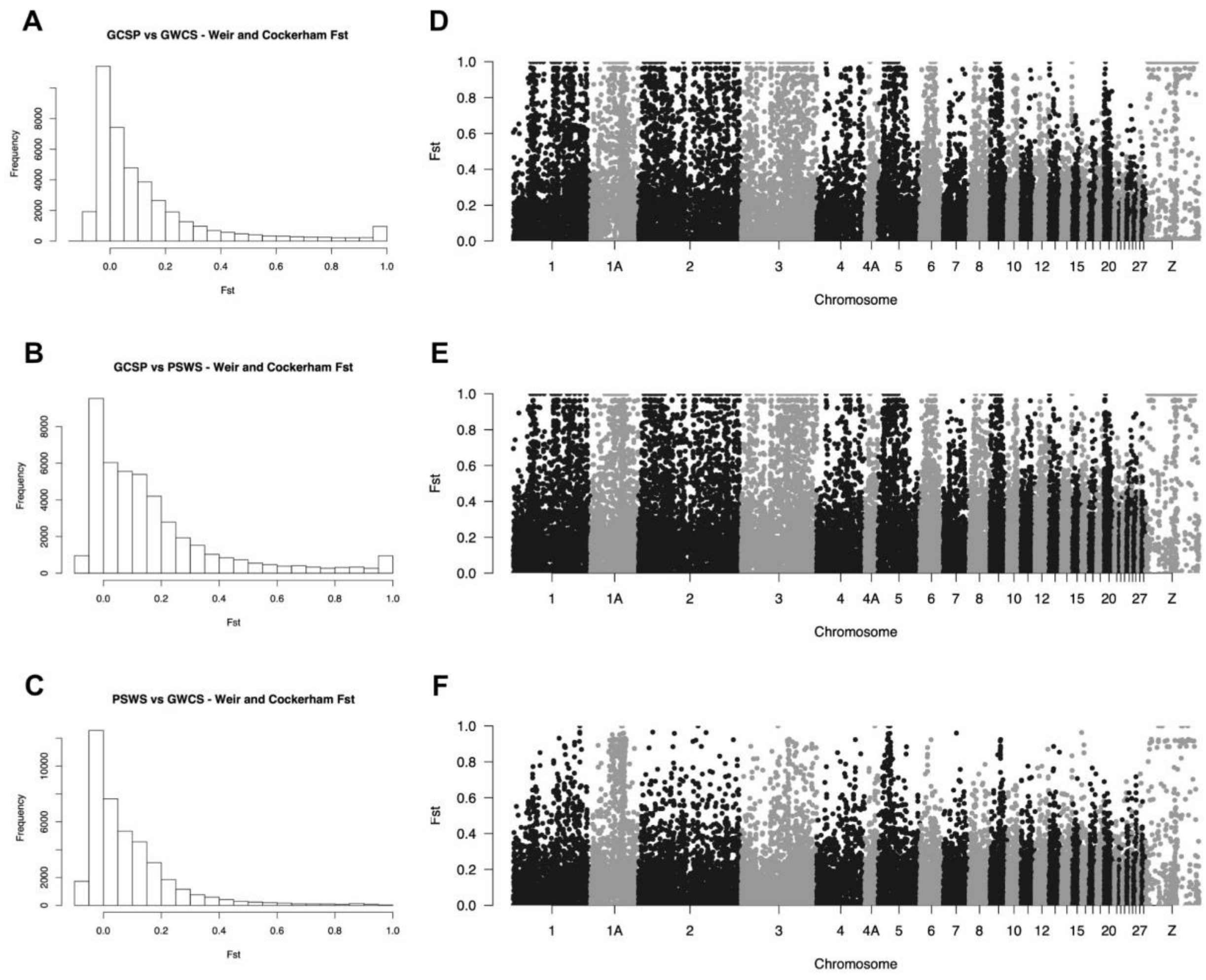

**Figure S5.** Summary of $F_{ST}$ at each biallelic variant SNP. (A-C) Histograms showing the distribution of $F_{ST}$ values for *Z. atricapilla* versus *Z. l. gambelii* (A), *Z. atricapilla* versus *Z. l. pugetensis* (B), and *Z. l. gambelii* versus *Z. l. pugetensis* (C). (D-F) Manhattan plots showing $F_{ST}$ at each SNP against the genomic position for *Z. atricapilla* versus *Z. l. gambelii* (D), *Z. atricapilla* versus *Z. l. pugetensis* (E), and *Z. l. gambelii* versus *Z. l. pugetensis* (F).

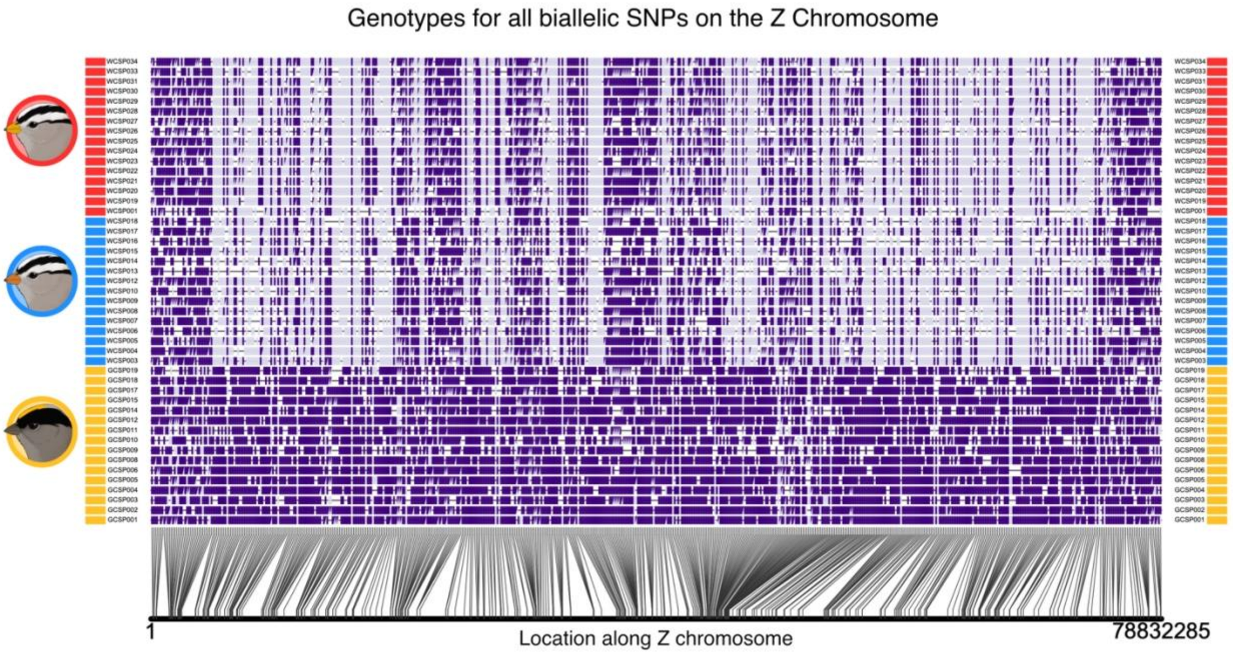

**Figure S6.** Genotypes at all biallelic SNPs on the Z chromosome. The most common genotype in *Z. atricapilla* is plotted in dark purple, and the alternative allele is plotted in light purple. Each row represents an individual in the dataset, and rectangles next to rows represent the taxon of that individual, with *Z. atricapilla* in yellow, *Z. l. gambelii* in blue, and *Z. l. pugetensis* in red.

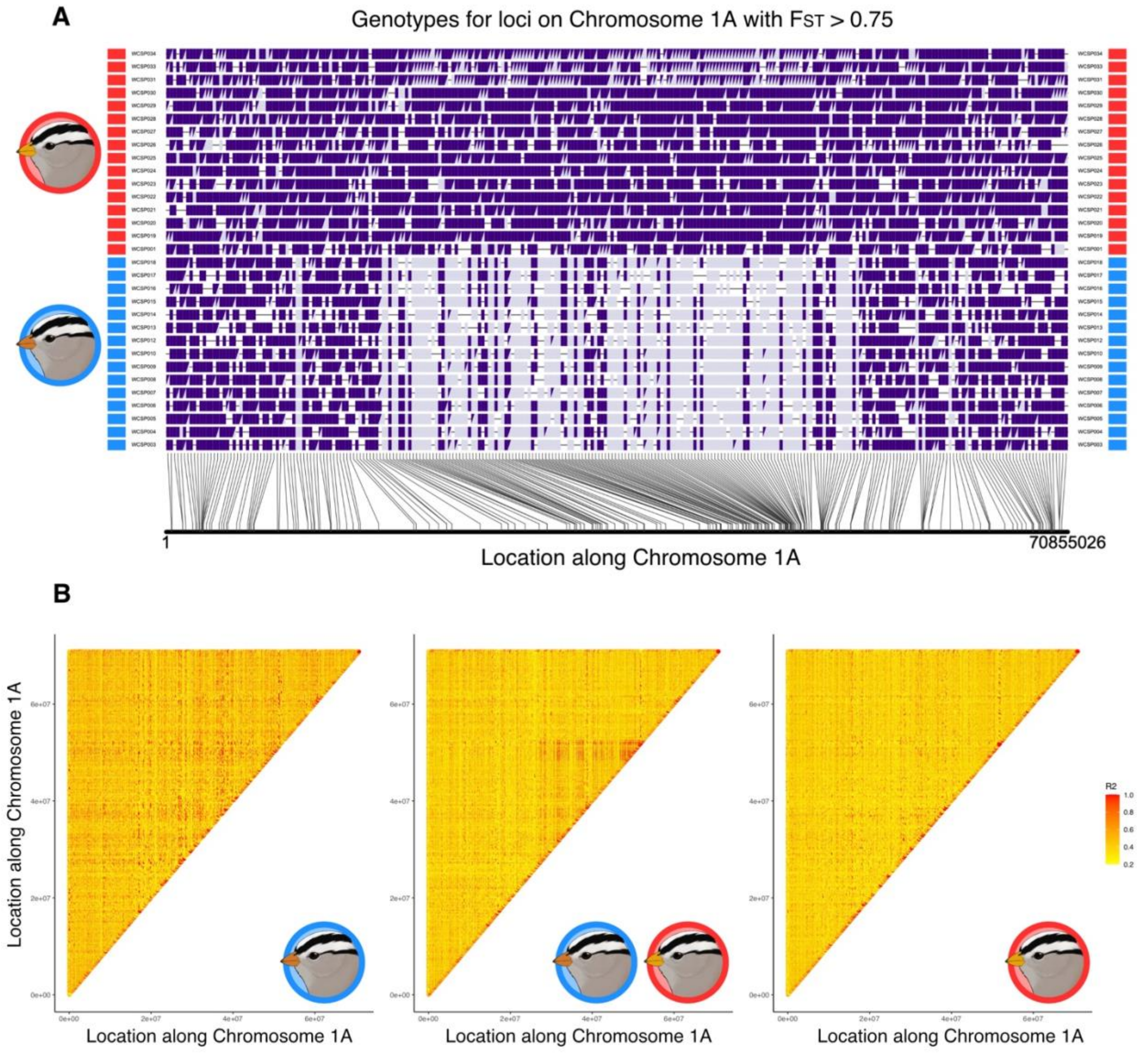

**Figure S7** Equivocal evidence for an inversion on chromosome 1A. (A) Genotypes at all biallelic SNPs that show an $F_{ST}$ > 0.75 on chromosome 1A. The most common genotype in *Z. l. pugetensis* is plotted in dark purple, and the alternative allele is plotted in grey. Each row represents an individual in the dataset, and rectangles next to rows represent the taxon of that individual, with *Z. l. gambelii* in blue, and *Z. l. pugetensis* in red. (B) Heatmaps of linkage disequilibrium (LD) across the Z chromosome when calculated between just *Z. l. gambelii* (left), both groups (middle), and just *Z. l. pugetensis* (right).

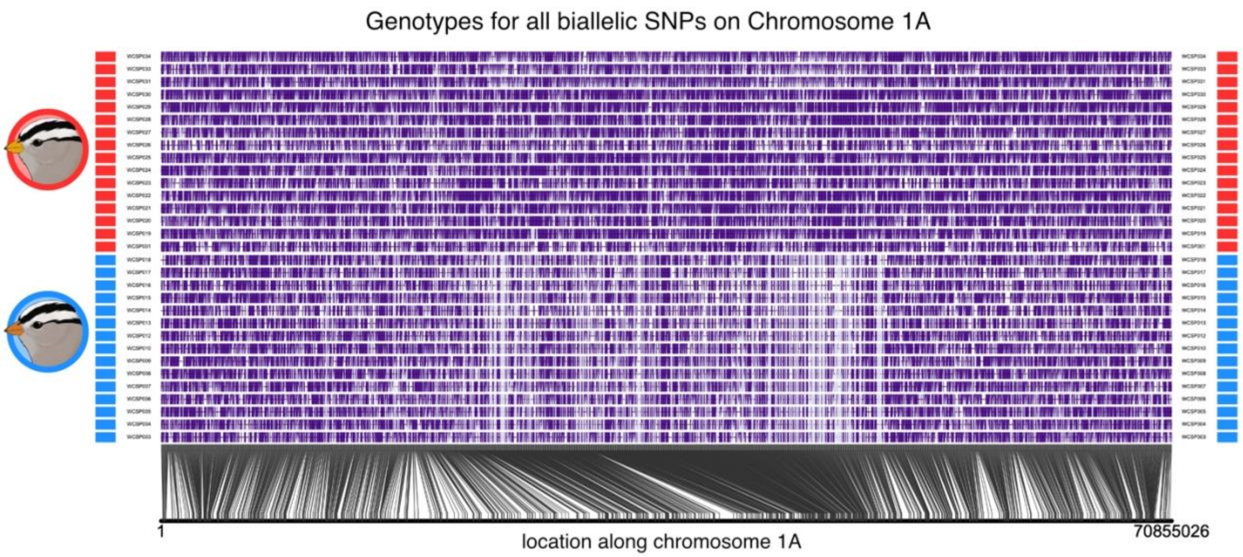

**Figure S8.** Genotypes at all biallelic SNPs on chromosome 1A. The most common genotype in *Z. l. pugetensis* is plotted in dark purple, and the alternative allele is plotted in light purple. Each row represents an individual in the dataset, and rectangles next to rows represent the taxon of that individual, with *Z. l. gambelii* in blue and *Z. l. pugetensis* in red.
